# Single-cell map of the healthy human immune system across the lifespan reveals unique infant immune signatures

**DOI:** 10.1101/2025.07.28.667181

**Authors:** Djamel Nehar-Belaid, Asa Thibodeau, Alper Eroglu, Radu Marches, Giray Eryilmaz, Derya Unutmaz, Chris P. Verschoor, Jinghua Gu, Uthra Balaji, Asunción Mejías, Virginia Pascual, George A. Kuchel, Octavio Ramilo, Jacques F. Banchereau, Duygu Ucar

**Affiliations:** The Jackson Laboratory for Genomic Medicine, Farmington, CT 06032 USA; Department of Cell and Molecular Biology, Karolinska Institute, Stockholm, 171 77, Sweden; SciLifeLab; Solna, Stockholm, 171 65, Sweden; Health Sciences North Research Institute; NOSM University; Sudbury, ON P3E 2H3, Canada; Department of Medicine, McMaster University; Hamilton, ON L8S 4L8 Canada; Department of Infectious Diseases, St. Jude Children’s Research Hospital, 262 Danny Thomas Place, Memphis, TN 38105; Drukier Institute for Children’s Health and Department of Pediatrics, Weill Cornell Medicine, New York, NY; UConn Center on Aging, University of Connecticut, Farmington, CT, USA; Immunoledge LLC, Montclair, NJ, USA; Institute for Systems Genomics, University of Connecticut Health Center, Farmington, CT, USA

**Author notes:** Co-senior authors (George A. Kuchel, Octavio Ramilo, Jacques F. Banchereau, Duygu Ucar).

## Abstract

The human immune system undergoes continuous remodeling from infancy through old age, yet the timing and trajectory of these changes across the lifespan remain poorly defined. To address this, we profiled peripheral blood mononuclear cells from 95 healthy individuals (ages 2 months to 88 years), including infants (n=27), children (n=23), adults (n=18), and older adults (n=27) using scRNA-seq and snATAC-seq. MAIT and γδ T cells showed a “Rise and fall” pattern, which rise in childhood, peak in young adulthood, and decline with age. CD8^+^ T cells were the most affected by aging with decreasing naïve T cells and increasing GzK^+^ CD8^+^ T cells and TEMRA cells. Infants had lower myeloid/lymphoid ratio, with a distinct composition marked by increased frequencies of CD16^+^ monocytes and plasmacytoid dendritic cells and reduced frequencies of CD14^+^ monocytes and conventional DCs. Their adaptive immune compartment also displayed unique features, including constitutive interferon-stimulated gene expression in T and B cells, and an expanded SOX4^+^ populations in naïve CD4^+^, naïve CD8^+^ and γδ T cells, comprising ∼30% of the naïve T cell pool. SOX4^+^ naïve CD4^+^ T cells displayed a Th2 epigenetic signature. This map provides critical insights into human immune system dynamics across the lifespan, emphasizing unique features of the infant immune system.

## Main

The human immune system is highly dynamic, continuously shaped by environmental exposures, infections, and vaccinations across the lifespan. Immune development begins in utero and changes are especially pronounced during early life and older age. In infancy, the immune system undergoes rapid transitions as maternal antibodies wane and adaptive immunity develops^1,2,3^. In contrast, aging is associated with accumulating of memory T cells, a decline in naïve cell populations, and a chronic/persistent low-grade inflammation^4–6^. Understanding how immunity develops, evolves and declines across the lifespan is essential for designing age-appropriate therapies and vaccines for both infectious and non-infectious age-associated diseases.

Recent advances in high-throughput genomics, particularly single-cell technologies, have revolutionized our ability to dissect the cellular and molecular architecture of the human immune system. Studies using single-cell RNA sequencing (scRNA-seq) of peripheral blood mononuclear cells (PBMCs) from young and older adults have identified age-associated immune features, such as expansions of HLA-DR⁺ CD4⁺ memory T cells and GZMK⁺ CD8⁺ T cells linked to chronic inflammation^7,8^. Studies of centenarians have revealed cell-type-specific signatures of exceptional longevity, such as upregulation of DNA damage response pathways^9^. While there are many studies focusing on immune aging, very few studies have focused on early life, especially the first months of infancy. This is due to logistical and practical challenges of recruiting and collecting samples in this population.

Infancy represents a uniquely critical window for immune development as initial exposures to certain antigens and priming occur and have direct implications for pediatric and overall immune health. Due to their immature immune system, infants are more vulnerable to severe infections from respiratory pathogens like influenza and respiratory syncytial virus (RSV), while paradoxically showing resilience against other viral respiratory infections such as SARS-CoV-2 that disproportionately affect older adults^10,11,12,13^. These contrasts suggest that infant immunity is qualitatively distinct and may also offer insights into protective mechanisms against certain pathogens that decline with age.

In this study, we leveraged single-cell RNA and single-nucleus ATAC sequencing to profile PBMCs from 95 healthy individuals encompassing the human lifespan, from 2-month-old infants to 88-year-old adults. This dataset allowed us to map immune development, maturation, and aging with an unprecedented resolution. Our analyses identified infancy as the most dynamic period of immune remodeling with unique features, particularly a predominance of plasmacytoid dendritic cells (pDCs), an expanded pool of SOX4⁺ naïve T cells, and constitutive interferon signaling in adaptive lymphocytes. These findings highlight early-life immune features, some of which might serve as novel targets to better support infant immunity and may also inform strategies to enhance immune resilience in older adults.

## Results

### Infant PBMCs stand apart from those of children and adults

We profiled peripheral blood mononuclear cells (PBMCs) from a cohort of 95 healthy individuals (52 females, 43 males; **Extended Data Fig. 1a**) spanning ages 2 months to 88 years (**Supplementary Table 1**) using single-cell RNA-sequencing (scRNA-seq). 581,724 cells were profiled across four age groups: infants (n=27, 2-18 months, 182,164 cells), children (n=23, 11-17 years, 138,973 cells), young adults (n=18, 22-38 years, 125,811 cells), and older adults (n=27, 65-88 years, 134,776 cells) (**Fig. 1a, Extended Data Fig. 1b,c and Supplementary Table 2**). We identified nine major immune cell subsets based on canonical marker gene expression: CD4^+^ T, CD8^+^ T, γδ T cells, B cells, NK cells, monocytes, dendritic cells (DCs), plasma cells (PCs), and hematopoietic stem and progenitor cell (HSPC) (**Fig. 1b,c and Supplementary Table 3a**). Our analysis included both frequencies within PBMCs and frequencies within a lineage.

**Fig. 1.**
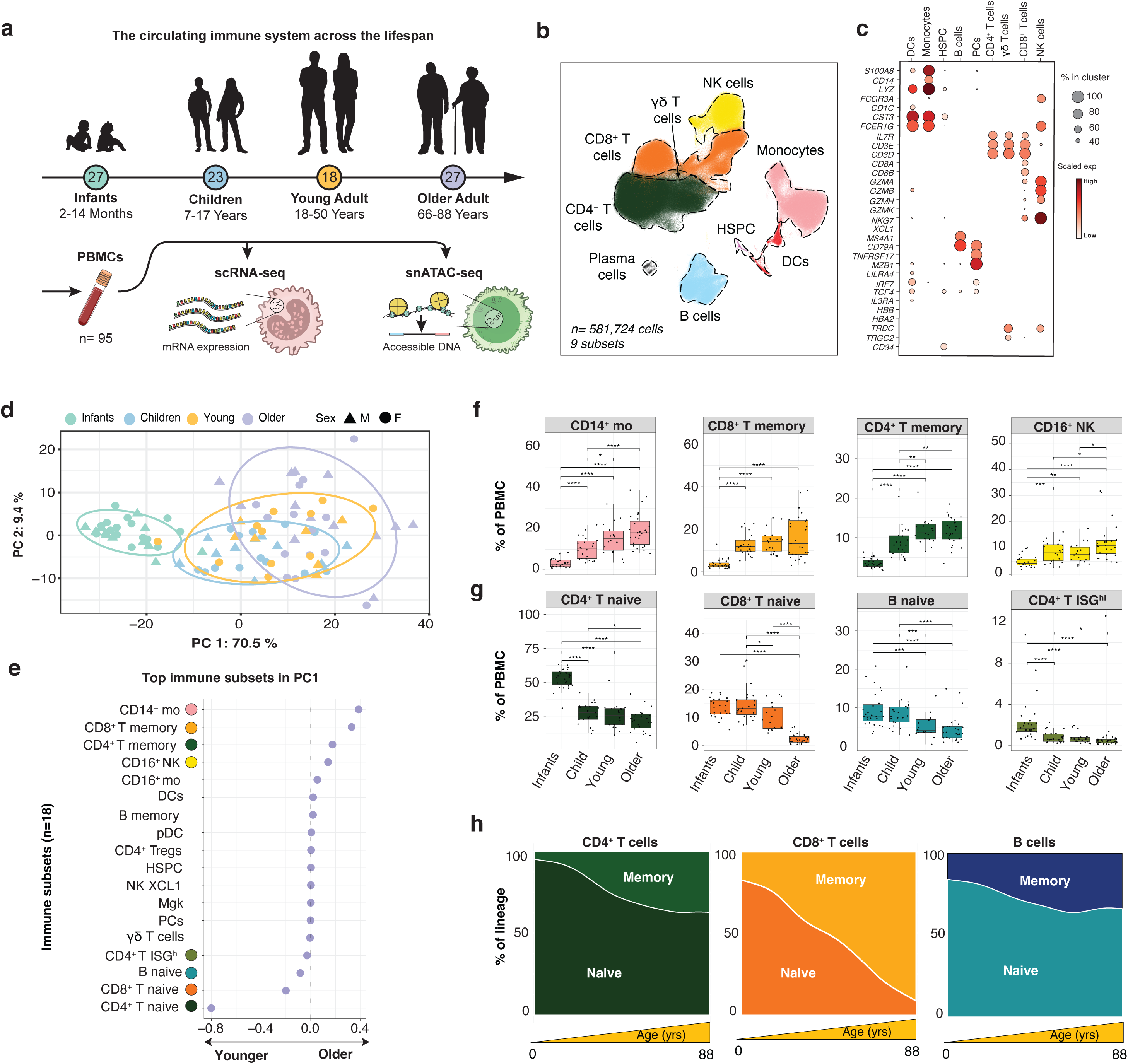
Study design and PBMC scRNA-seq data. **a**, Study design. **b**, UMAP plot representing 581,724 cells colored by immune lineages (n=9). **c**, Dot plot showing the expression values of selected genes (x-axis) across each cluster (y-axis). Dot size represents the percentage of cells expressing the marker of interest. Color intensity indicates the mean expression within expressing cells. **d**, Principal Component Analysis based on cell compositional frequencies among the 95 donors. **e**, Immune subsets contributing to PC1. **f**, Boxplots for the cell frequencies for top four subsets discriminating infants based on PC1 loadings. P values were calculated using a two-sided t-test comparing the mean within each age group. *, P<0.05; **, P<0.01; ***, P<0.001: ****, P<0.0001, and ns: non-significant. The upper and lower bounds represent the 75% and 25% percentiles, respectively. **g**, Boxplots for cell frequencies discriminating other age groups, based on PC1 loadings. **h**, Shifts from naïve to memory cells with age for CD4^+^ T, CD8^+^ T, and B cells.

Both adaptive and innate immune cells exhibited age-related remodeling (**Extended Data Fig. 2a**). Innate immune cell frequencies increased with age, including monocytes (R=0.69; p<0.001), DCs (R=0.55; p<0.001), and NK cells (R= 0.49; p<0.001). Conversely, adaptive immune cell frequencies declined with age, including CD4^+^ T cells (R=-0.54; p<0.001), B cells (R=-0.42; p<0.001), plasma cells (R=-0.3; p=0.0041), and γδ T cells (R=-0.58, p<0.001). Notably, total CD8^+^ T cell frequencies remained stable throughout lifespan (R=-0.069; p=0.51) (**Extended Data Fig. 2a**).

We further clustered PBMCs into 18 immune subsets identifying CD14^+^ and CD16^+^ monocytes, CD16^+^ and XCL1^+^ NK cells, naïve and memory T and B cells, interferon-stimulated gene-expressing ISG^hi^ CD4^+^ T cells, and Tregs (**Extended Data Fig. 2b,c and Supplementary Table 3b**). Principal Component Analysis (PCA) on the frequency of these 18 immune subsets separated infants from other age groups, with PC1 explaining 70% of the variance (**Fig. 1d**). This separation was largely driven by age-associated frequencies of naïve CD4^+^ T cells (∼53% of infant PBMCs *vs.* ∼22% in older adults) and low frequency of CD14^+^ monocytes (∼3% of infant PBMCs *vs.* ∼19% in older adults) (**Fig. 1e**). We also detected ISG^hi^ CD4^+^ T cells that constitutively express interferon-stimulated genes (*e.g., ISG15*). ISG^hi^ CD4^+^ T cells were more abundant in infants, as ∼2.2% of their PBMCs were ISG^hi^, compared to ∼0.9% in adults (**Fig. 1f,g and Extended Data Fig. 2d,e**). Infant PBMCs were predominantly composed of naïve T and B lymphocyte. Naïve CD4^+^ T cells decreased sharply after infancy, naïve B cells showed more variation between children and adults, while naïve CD8^+^ T cells declined in older adults, suggesting that lymphocyte populations follow distinct kinetic patterns across different age groups (**Fig. 1g**).

Within their respective lineages, naïve CD4^+^ T and B cells declined gradually, with relatively modest changes between younger and older adults. In contrast, naïve CD8^+^ T cells showed a sharp and continuous decline across the lifespan (**Fig. 1h**).

These data revealed that infant’s immune system stands apart from the other age groups. The following sections will further dissect immune remodeling across the human lifespan in each lineage.

### Infants display low cDCs and high pDCs

To investigate myeloid cell dynamics, we analyzed DC subsets (n=6,617 cells): cDC1 (*CLEC9A*, *XCR1*), cDC2 (*CD1C*, *CLEC10A*), monocyte-derived DCs (moDCs; *CD14*), AXL^+^ DCs (*AXL*), and pDC (*IRF7*) (**Fig. 2a, Extended Data Fig. 3a and Supplementary Table 4a**). DC frequencies increased with age, including cDC1 (R=0.4; p<0.001), cDC2 (R=0.56; p<0.001), moDCs (R=0.56, p<0.001), AXL^+^ DC (R=0.32, p=0.011), and pDC (R=0.29, p=0.0041) (**Extended Data Fig. 3b**). The largest expansion in DCs occurred from infancy to childhood, with moDC and cDC2 populations expanding significantly during this time (p<0.001) (**Fig. 2b and Extended Data Fig. 3b**). In contrast, cDC1 frequencies increased during transition from childhood to adulthood (p=0.012). DC subset frequencies remained stable between young and older adults, indicating that most DC remodeling happens earlier in life.

**Fig. 2.**
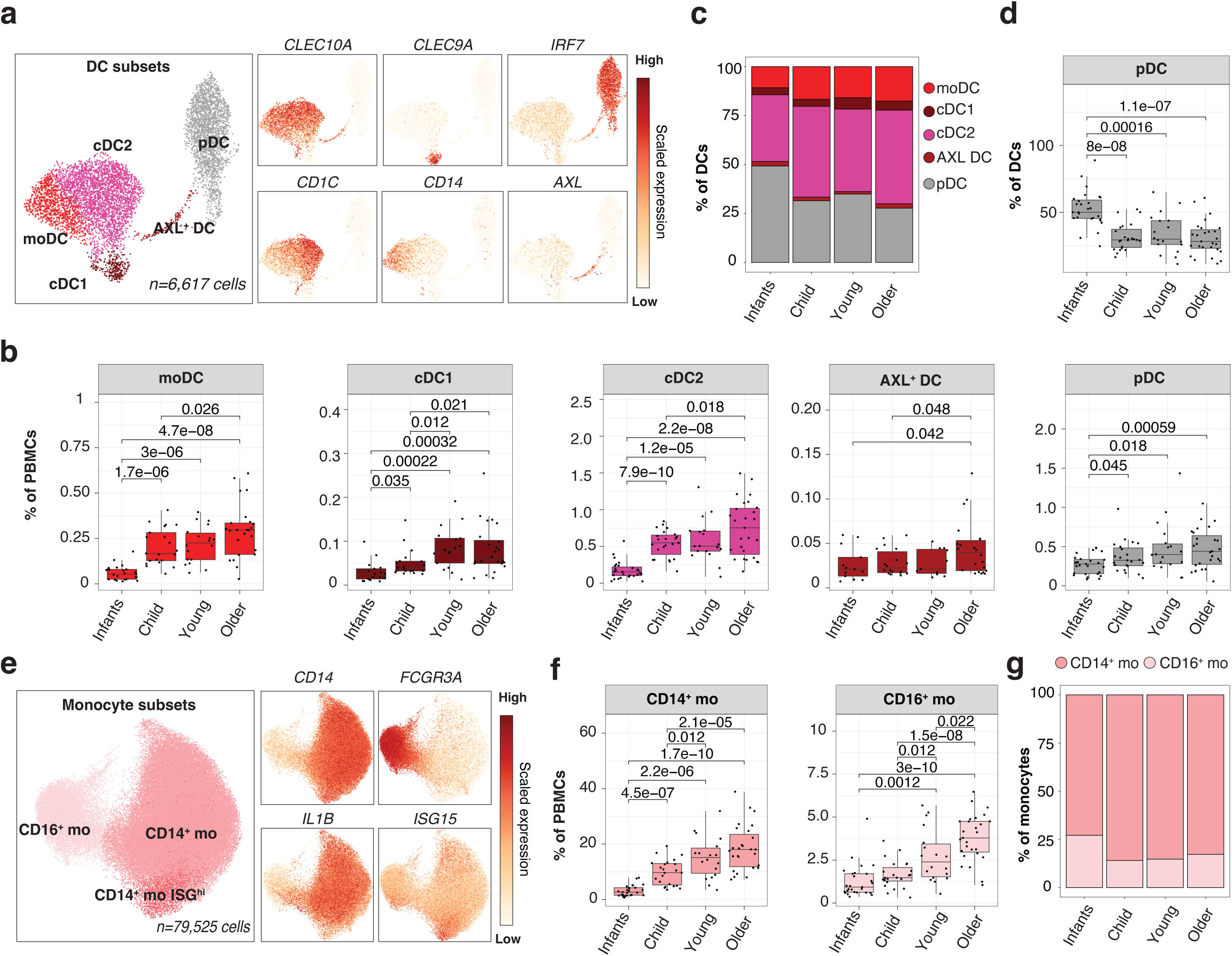
Changes in DCs and monocytes across lifespan. **a**, UMAP plot representing DC subsets (n=6,617) along with feature plots for marker gene. **b**, Boxplots comparing the frequencies of five DC subsets in PBMCs for 95 individuals categorized into four age groups. P values were calculated using a two-sided t-test comparing the mean within each age group. The upper and lower bounds represent the 75% and 25% percentiles, respectively. **c**, Barplots comparing the distribution of DC subsets within DCs across age groups. **d**, Boxplot showing distribution of pDCs within DCs across age groups. **e**, UMAP plot representing 79,525 monocytes colored by subset along with feature plots for marker gene. **f**, Boxplots comparing the frequencies of CD14^+^ and CD16^+^ monocytes in PBMCs for 95 individuals categorized into four age groups. P values were calculated using a two-sided t-test comparing the mean within each age group. The upper and lower bounds represent the 75% and 25% percentiles, respectively. **g**, Barplots showing distribution of monocyte subsets within monocytes across age groups.

Although infants had fewer DCs overall, their DC composition was distinct from other age groups. Infants exhibited a predominance of pDCs (∼50% of total DCs), while older adults had higher frequencies of cDC2 (∼48% of DCs) and moDCs (∼18% of DCs) (**Fig. 2c,d and Extended Data Fig. 3c**).

Monocyte analysis identified three subsets: CD14^+^ monocytes (*CD14;* n=63,806), CD16^+^ monocytes (*FCGR3A;* n=12,918), and CD14^+^ monocytes expressing interferon-stimulated genes (ISG^hi^; *ISG15*; n= 2,801) (**Fig. 2e, Extended Data Fig. 3d and Supplementary Table 4b**). Both CD14^+^ and CD16^+^ monocyte frequencies increased with age, with CD16^+^ monocytes exhibiting a more pronounced expansion (p=0.022) between younger and older adults (**Fig. 2f,g**).

Infants had overall fewer monocytes than older age groups, and their composition was distinct. Specifically, infants had a higher frequency of CD16^+^ monocytes (∼26% of total monocytes versus ∼15% in other age groups) and fewer CD14^+^ monocytes (70% of total monocytes versus ∼81% in other age groups) (**Extended Data Fig. 3e-g**). Myeloid cell subsets frequencies did not change during the first 20 months of life (**Extended Data Fig. 3h-k**).

### Infants exhibit higher frequencies of immature and proliferating NK cells

NK cells (n=52,378 cells) clustered into four subsets: CD16^+^ NK (*FCGR3A*), KLRC2^+^ CD16^+^ NK (*KLRC2*/NKG2C), CD56^bright^ NK (*XCL1*, *GZMK*), and proliferating NK cells (proliferative, *MKI67*) (**Fig. 3a**). Mature CD16^+^ NK cell frequencies increased with age (R=0.51, p<0.001), constituting ∼4.5% of PBMCs in infants and increasing to ∼11% in older adults (**Fig. 3b, Extended Data Fig. 4a and Supplementary Table 4c)**. The frequencies of other NK subsets remained largely unchanged across age groups, but donor-specific expansion of KLRC2^+^ CD16^+^ NK cells (adaptive NK cells) was observed (**Fig. 3b**). Infants exhibited a distinct NK cell profile; characterized by higher frequencies of immature CD56^bright^ and proliferating NK cells. Mature CD16^+^ NK cells increased with age (**Fig. 3c and Extended Data Fig. 4b**). Proliferating NK cell frequency declined during the first months of life (R=-0.46, p=0.021) (**Fig. 3d and Extended Data Fig. 4c**).

**Fig. 3.**
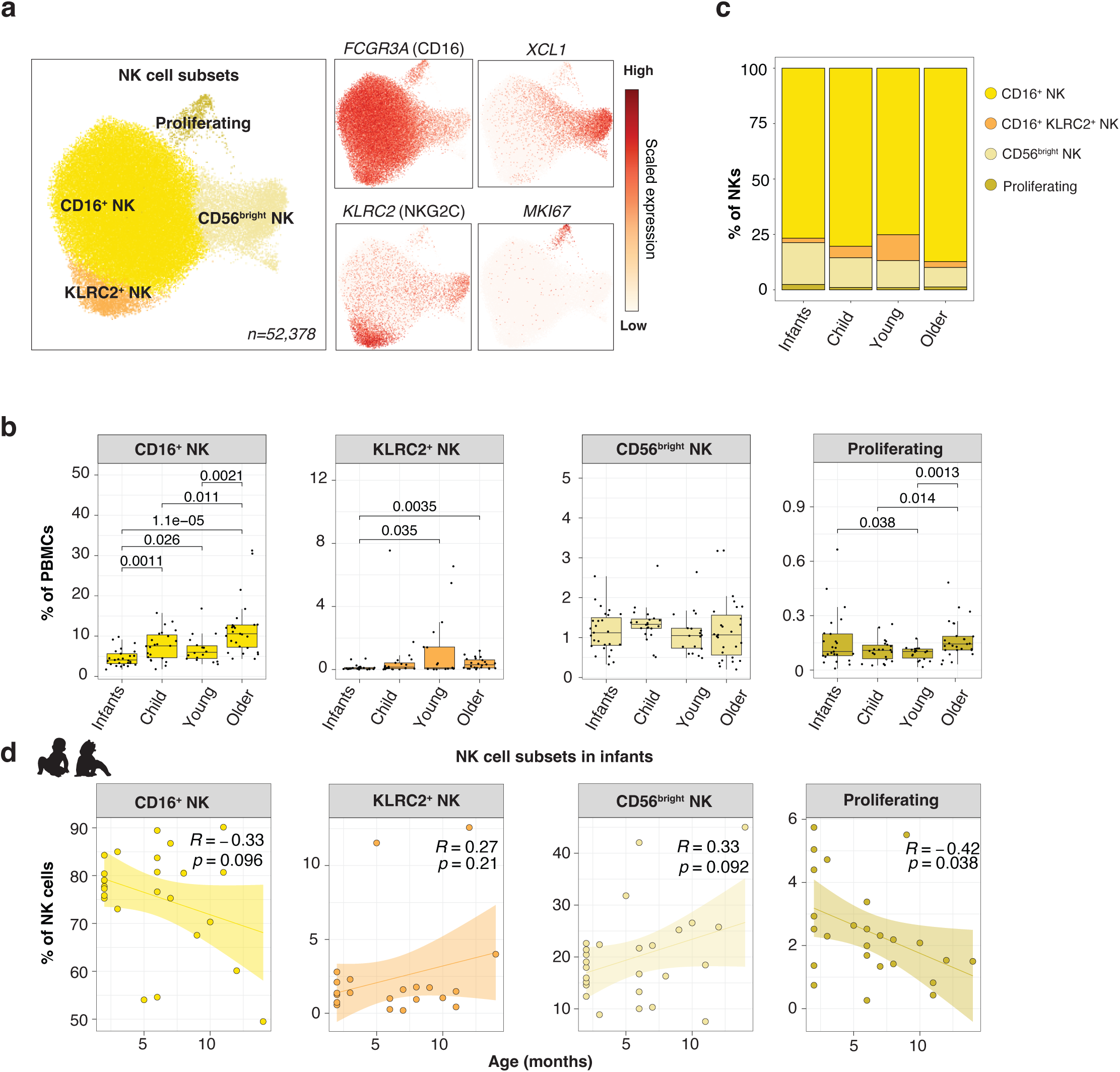
Changes in NK cells across lifespan. **a**, UMAP plot representing NK subsets (n=52,378) along with feature plots for marker gene. **b**, Boxplot comparing the frequencies of NK cell subsets in PBMCs for 95 individuals across four age groups. P values were calculated using a two-sided t-test comparing the mean within each age group. The upper and lower bounds represent the 75% and 25% percentiles, respectively. **c**, Barplots summarizing NK cell subsets within lineage for each age group. **d**, NK cell subset frequencies within lineage (y-axis) versus age of infants in months (x-axis). Each line represents the best-fitted linear regression, with the shading showing the 95% confidence intervals. R represents the Pearson correlation coefficient.

In summary, mature cytotoxic CD16^+^ NK cells progressively increased with aging, reflecting a shift toward NK cell maturation over the lifespan.

### Infant B cells are enriched in naïve, transitional, and ISG^hi^ subsets

B cells (n=55,242) clustered into six subsets: naïve (*IGHD*, *CCR7*), transitional (*CD9*, *MME*), memory (*CD27*), age-associated B cells (ABCs; *ITGAX*, *TBX21*, *FCRL5*, lack of *CD27* and *IGHD*), ISG^hi^ (*ISG15*, *IFI44*, *MX1*), and plasma cells (PCs; *JCHAIN*, *TNFRSF17*/BCMA) (**Fig. 4a and Extended Data Fig. 4d and Supplementary Table 4d**). Naïve and transitional B cell frequencies were higher in infants and children than in adults and negatively correlated with age (R=-0.32 and p<0.001 for naive; R=-0.5 and p<0.001 for transitional) (**Fig. 4b and Extended Data Fig. 4e**). Infants also had higher frequencies of ISG^hi^ B cells compared to young (p=0.034) and older adults (p=0.04) within their PBMCs (**Fig. 4b**). Infants and children had higher frequency of plasma cells (PC) compared to older adults (p=0.023 and p=0.0012, respectively) (**Fig. 4b**). Within B cells, ABC frequencies increased (R=0.42, p<0.001), while transitional B cells declined (R=-0.49, p<0.001) with age (**Fig. 4c and Extended Data Fig. 4f**). As shown in **Fig. 4d** focusing on infants; ABC frequencies increased rapidly (R=0.59, p=0.0018), while transitional B cell frequencies declined (R=-0.54, p=0.0038).

**Fig. 4.**
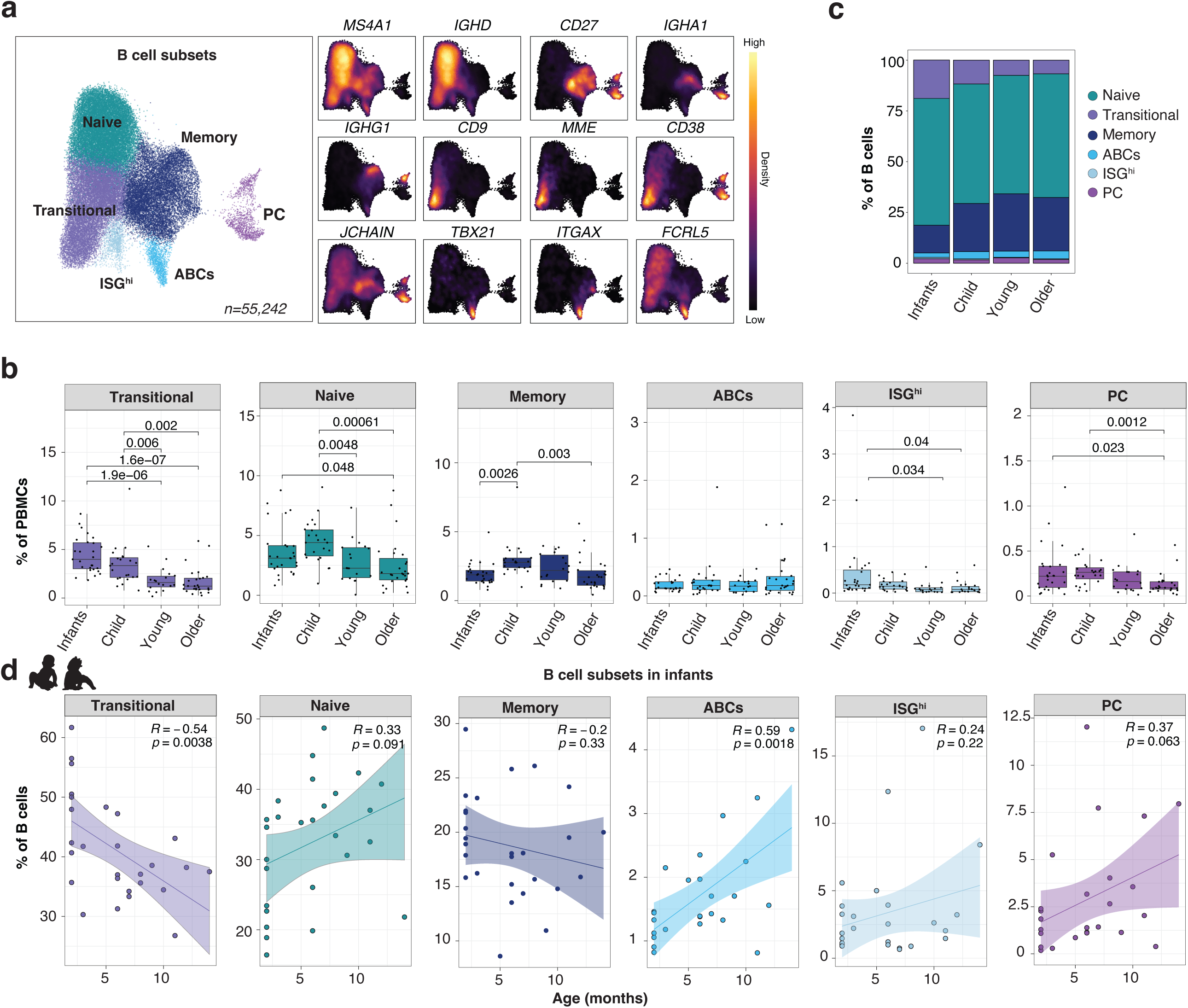
Changes in B cells across lifespan. **a**, UMAP plots representing B subsets (n=55,242; right) along with feature plots for marker gene (left). **b**, Boxplots comparing the frequencies of B cell subsets in PBMCs across 95 individuals across four age groups. P values were calculated using a two-sided t-test comparing the mean within each age group. The upper and lower bounds represent the 75% and 25% percentiles, respectively. **c**, Barplots summarizing B cell subsets within lineage for each age group. **d**, B cell subset frequencies within lineage (y-axis) versus age of infants in months (x-axis). Each line represents the best-fitted linear regression, with the shading showing the 95% confidence intervals. R represents the Pearson correlation coefficient.

Key features of infant B cells included: (i) increased frequencies of transitional B cells (∼40% of infant B cells), which declined in childhood (25%) and (ii) higher ISG^hi^ B cell frequencies in infants (∼4% of B cells) compared to other age groups (∼1.5% of B cells) (**Fig. 4c and Extended Data Fig. 4f**).

### HLA-DR^+^ CD4^+^ T cells, CD4^+^ TEMRAs, and memory Tregs expand with age

CD4^+^ T cells (259,600 cells) clustered into four major subsets: naïve (*CCR7*), naïve ISG^hi^ (*ISG15*), regulatory (Tregs; *FOXP3*), and memory (*S100A4*) CD4^+^ T cells (**Fig. 5a and Supplementary Table 4e**). Naïve CD4^+^ T cell frequencies declined with age (R=-0.63, p<0.001) (**Extended Data Fig. 5a**), representing >50% of infant PBMCs, 25% in children and ∼20% in young and older adults (**Fig. 5b**). ∼2.2% of infant’s PBMCs were naïve CD4^+^ T cells expressing ISGs in the absence of any reported acute infection or recent vaccination (**Supplementary Table 1**), compared with 0.9% in other age groups (**Fig. 5b**). Conversely, memory CD4^+^ T cells (R=0.6, p<0.001) and Treg (R=0.37, p<0.001) frequencies increased with age (**Extended Data Fig. 5a,b**). CD4^+^ T cell subsets from infants remained stable during development (**Extended Data Fig. 5c,d**).

**Fig. 5.**
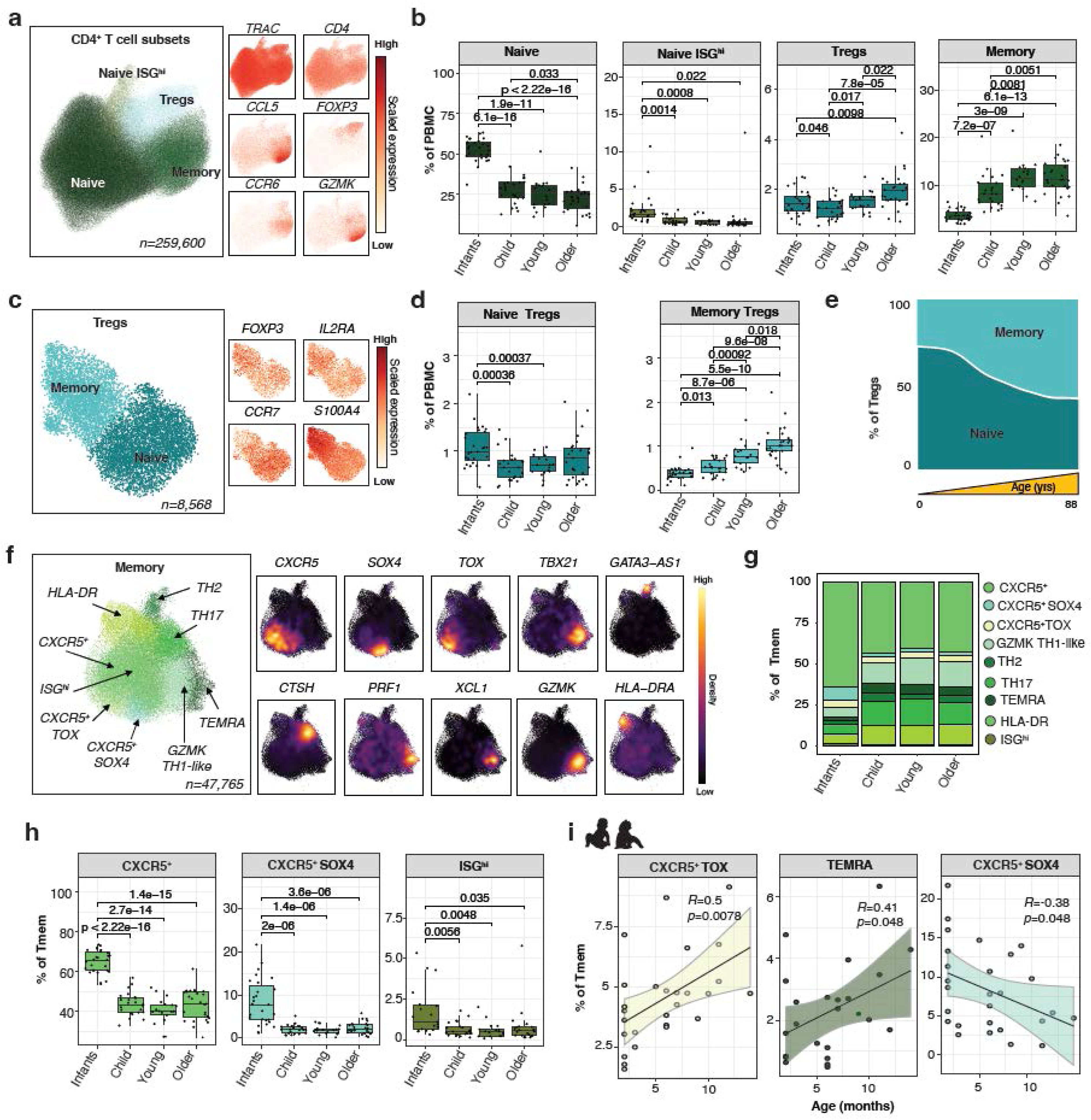
Changes in CD4^+^ T cells across lifespan. **a**, UMAP plot representing CD4^+^ T cell subsets (n=277,606) along with feature plots for marker genes. **b**, Boxplots comparing the frequencies of CD4^+^ T cell subsets in PBMCs across 95 individuals across four age groups. The upper and lower bounds represent the 75% and 25% percentiles, respectively. **c**, UMAP plot representing Treg subsets (n=8,568) and marker genes. **d**, Boxplots comparing the frequencies of Treg subsets in PBMCs across four age groups. The upper and lower bounds represent the 75% and 25% percentiles, respectively. **e**, Shifts from naïve to memory Treg subsets with age. **f**, UMAP plot representing CD4^+^ memory T cells subsets (n=47,765), with marker gene expression. **g**, Barplots comparing frequencies of CD4^+^ memory T cells within PBMCs across age groups. **h**, Boxplots comparing distribution of three subsets enriched in infants’ CD4^+^ memory T cells. The upper and lower bounds represent the 75% and 25% percentiles, respectively. **i**, CD4^+^ memory subset frequencies (y-axis) within lineage versus age of infants in months (x-axis). Each line represents the best-fitted linear regression, with the shading showing the 95% confidence intervals. R represents the Pearson correlation coefficient.

We further analyzed each of these compartments in detail (**Supplementary Table 4f**). Tregs were further clustered into naïve (*CCR7*) and memory (*S100A4*) subsets (**Fig. 5c**). Within PBMCs, naïve Tregs were more frequent in infants compared to children (p<0.001) and young adults (p<0.001) (**Fig. 5d**), whereas memory Treg frequencies increased with age (R=0.69, p<0.001) (**Extended Data Fig. 5e**). Within Tregs, the ratio of naïve to memory shifted from ∼75:25 in infants to 25:75 in older adults, marking a clear transition with aging (**Fig. 5e and Extended Data Fig. 5f**). Treg frequencies did not change significantly during infancy (**Extended Data Fig. 5g,h**).

CD4^+^ memory T cells represented 3% of infant PBMCs, ∼8% of children PBMCs and ∼11% of adult PBMCs (**Fig. 5b**). Memory CD4^+^ T cells clustered into nine subsets (**Fig. 5f**): CXCR5^+^ TFH-like (*CXCR5*), CXCR5^+^ TOX^+^ (*TOX*), CXCR5^+^ SOX4^+^ (*SOX4*), HLA-DR^+^ (*HLA-DRA*), TEMRA (*PRF1*, *NKG7*), TH1 (*GZMK*, *XCL1*), TH2 (*GATA3-AS1*), TH17 (*CTSH*), ISG^hi^ (*ISG15*) (**Extended Data Fig. 5i).** Most memory CD4^+^ subsets expanded with age, except for: CXCR5^+^ SOX4^+^ and ISG^hi^ (**Extended Data Fig. 5j & Extended Data Fig. 6a**).

Although infants had fewer memory CD4^+^ T cells, their memory compartment was uniquely enriched in three subsets: CXCR5^+^ TFH-like, CXCR5^+^ SOX4^+^, and ISG^hi^ cells (**Fig. 5g,h**). CXCR5^+^ TFH-like cells constituted ∼65% of infant CD4^+^ memory T cells, which rapidly declined to ∼40% in childhood. CXCR5^+^ SOX4⁺ T cells, a population that declined with age (R=-0.41, p<0.001), represented ∼9% of memory cells in infants (**Extended Data Fig. 6b**). Finally, ISG^hi^ CD4^+^ memory T cells were more abundant in infants compared to other age groups (1.2% *vs.* 0.55%) (**Fig. 5h**). Dynamics of memory CD4^+^ T cells during infancy revealed rapid increases in CXCR5^+^ TOX^+^ (R=0.5, p=0.0078) and TEMRA cells (R=0.41, p=0.048) (**Fig. 5i and Extended Data Fig. 6c,d**).

Memory CD4^+^ T cells were dynamically remodeled across lifespan, shifting from naïve to memory subsets, with infants exhibiting unique features: (i) high abundance of naïve CD4^+^ and naïve Tregs and (ii) low abundance of memory T cell populations, with an enrichment of three memory subsets, including CXCR5^+^ TFH-like cells.

### Rise and fall of MAIT and γδ T cells across the lifespan

CD8^+^ T cells (n=123,872 cells) clustered into seven subsets: naïve (*CCR7*), GZMK^+^ (*GZMK*), TEMRA (*GZMB, PRF1*), γδ (*TRDC*), MAIT (*ZBTB16*), and proliferating cells (*MIKI67*) (**Fig. 6a, Extended Data Fig. 7a and Supplementary Table 4g**). All CD8^+^ T subsets showed significant age-associated changes (**Fig. 6b**). Naive CD8^+^ T cell frequencies declined sharply with age (R=-0.76, p<0.001), constituting ∼78% of infant CD8⁺ T cells but only ∼17% in older adults (**Fig. 6c and Extended Data Fig. 7b,c)**. Frequencies of GZMK^+^ (R=0.56, p<0.001) and TEMRA cells (R=0.5, p<0.001) increased with age (**Fig. 6b-c and Extended Data Fig. 7b,c**), together comprising 75% of older adults’ CD8⁺ T cells (34% and 41%, respectively) compared to ∼15% in infants. The expansion of these memory CD8^+^ subsets began during infancy (**Extended Data Fig. 7d,e**).

**Fig. 6.**
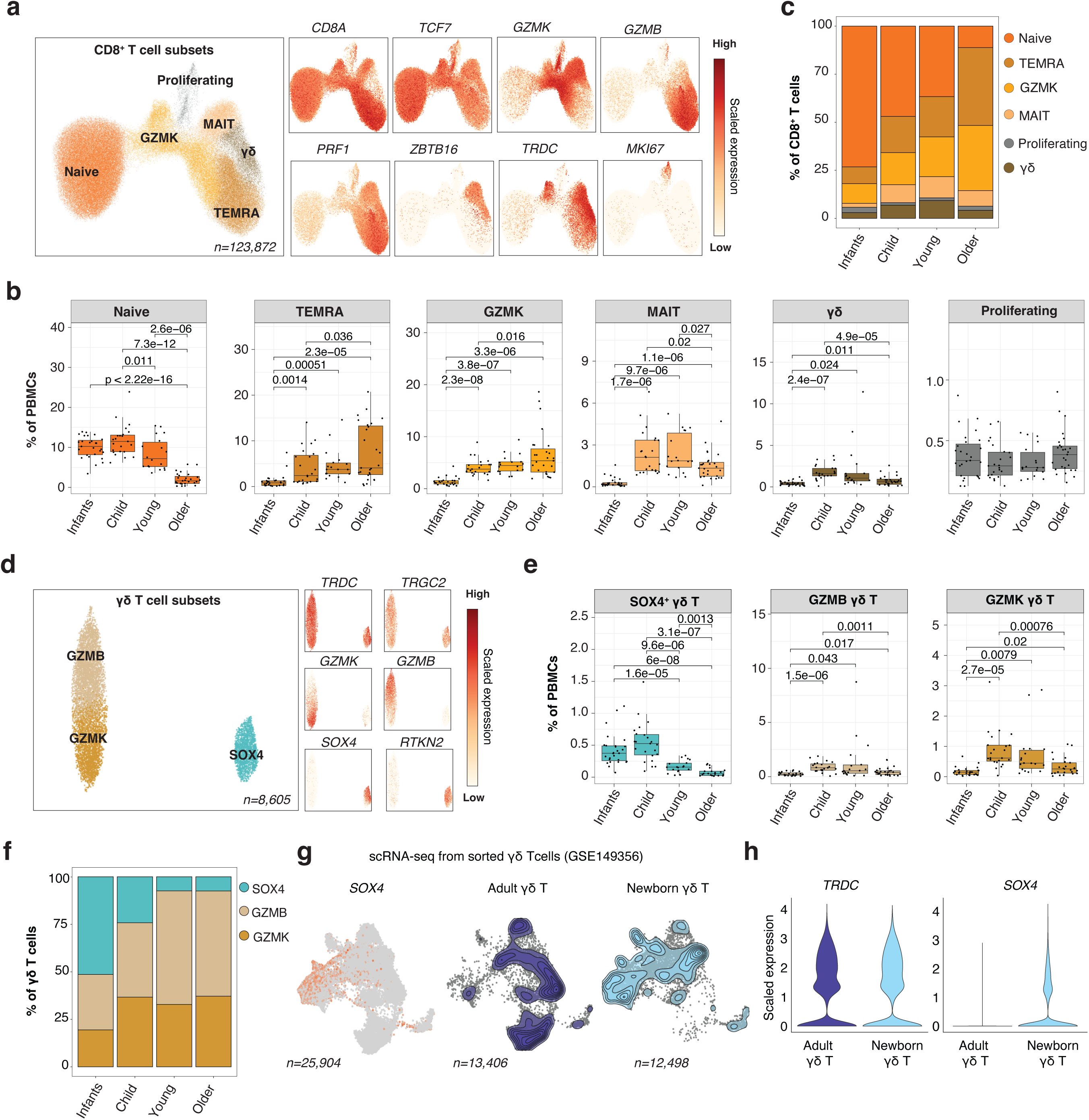
Changes in CD8^+^ T cells across lifespan. **a**, UMAP plot representing CD8^+^ T cell subsets (n=123,872), along with feature plots for marker gene. **b**, Boxplots comparing the frequencies of CD8^+^ T cell subsets in PBMCs across 95 individuals across age groups. The upper and lower bounds represent the 75% and 25% percentiles, respectively. **c**, Barplots showing distribution of CD8^+^ subsets within lineage across age groups. **d**, UMAP plot representing γδ T cell subsets and marker genes. **e**, γδ T cell subset frequencies in PBMCs across age groups. **f**, Bar plots showing distribution of γδ T cell subsets within lineage across age groups **g**, Left: SOX4 expression in public γδ T cell scRNA-seq data from adults and newborns (GSE149356). Right: The density of cells in adults and newborns. **h**, Violin plots showing the expression of *TRDC* and *SOX4* in γδ T cells from adults and newborns.

MAIT and γδ CD8^+^ T cell exhibited a “rise with maturation and fall with aging” pattern, as frequencies peaked in childhood and young adulthood before declining in older adulthood (**Fig. 6b**). MAIT cell frequencies were highest in children and young adults (up to 10% of PBMCs), but were rare in infants (<1%) (**Fig. 6b and Extended Data Fig. 7c**) and lower in older adults. γδ T cells clustered into three subsets (**Fig. 6d**): GZMB^+^, GZMK^+^, and naïve SOX4^+^ γδ T cells (**Supplementary Table 4h)**. All three subsets followed the “rise and fall” pattern (**Fig. 6e**), but a significant proportion of infant γδ T cells were SOX4^+^ (>50%) compared to other age groups (25% in children, <10% in adults) **(Fig. 6e,f and Extended Data Fig. 7f,g**). Reanalysis of publicly available sorted γδ T cell scRNA-seq data from newborns and adults^14^ confirmed high expression of *SOX4* in newborn γδ T cells (**Fig. 6g,h**).

CD8^+^ T cells significantly remodeled across lifespan. Infant CD8^+^ T cells were mostly composed of naïve CD8^+^ T cells. MAIT and γδ T cells exhibited a ‘rise and fall’ pattern. Infants γδ T cells were enriched in SOX4⁺ γδ T cells. As expected, there was a progressive expansion of memory subsets with age.

### SOX4^+^ naïve T cells enriched in infants exhibit stemness features

20% of both naïve CD4^+^ and CD8^+^ T cells highly expressed SOX4 (**Fig. 7a,b**). These SOX4^+^ T cells were predominantly found in infants, representing ∼18.5% of their PBMCs, compared with <3% in older adults (**Fig. 7c,d, Extended Data Fig. 8a-d**). The frequency of SOX4^+^ naïve T cells declined rapidly during infancy (**Fig. 7e,f and Extended Data Fig. 8e,f**). In contrast, SOX4^−^ naïve T cells exhibited a more gradual decline that started later during childhood.

**Fig. 7.**
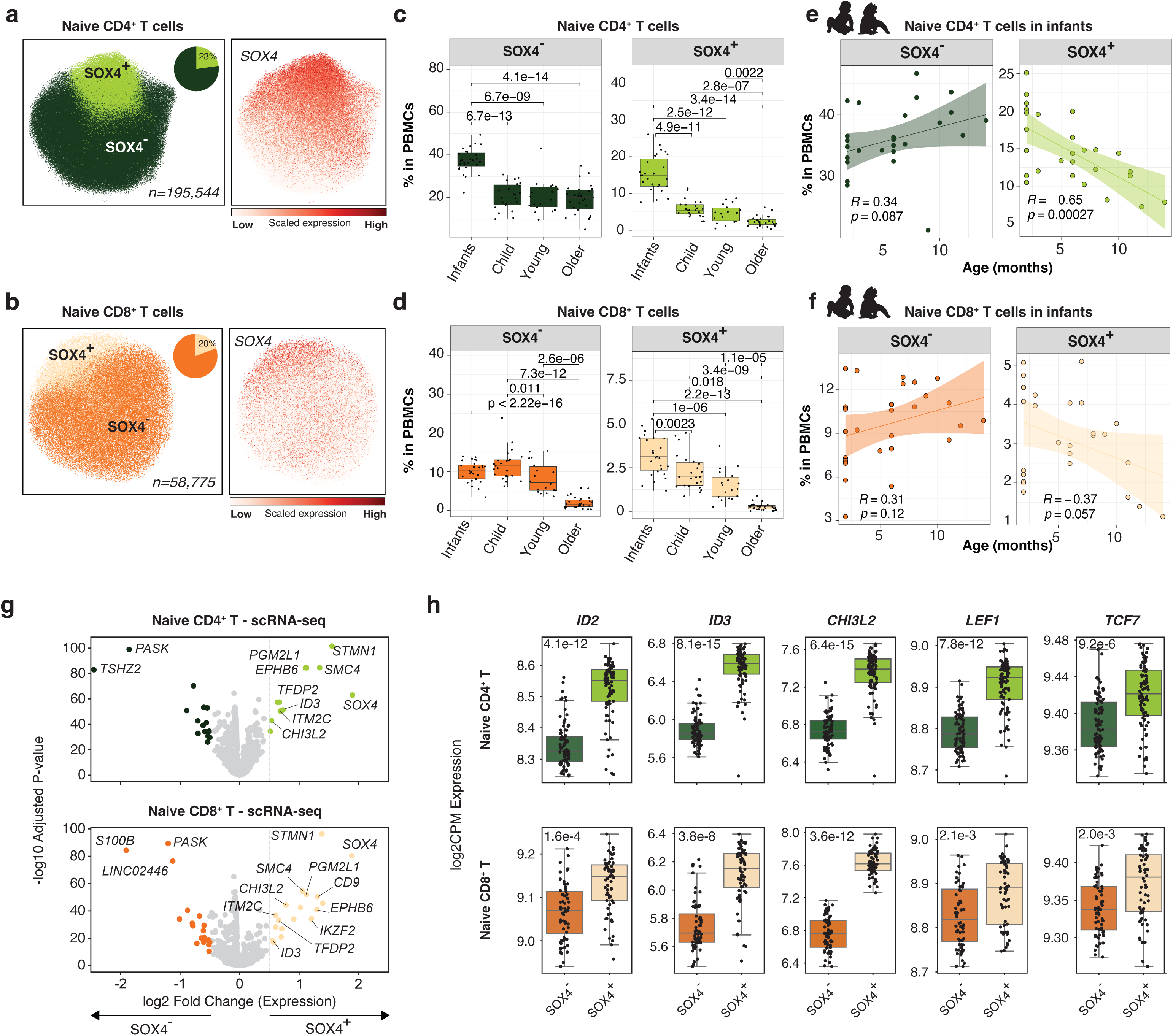
Abundance of SOX4^+^ T cells in infants. **a**, UMAP plot representing naive CD4^+^ T cell subsets (n=195,544; left) and SOX4 gene expression (right). **b**, UMAP plot representing naive CD8^+^ T cell subsets (n= 58,775; left) and SOX4 gene expression (right). **c**, Boxplots comparing the frequencies of SOX4^+^ and SOX4^−^ naïve CD4^+^ T cell subsets and naïve CD8^+^ T cells (**d**) in PBMCs across four age groups. The upper and lower bounds represent the 75% and 25% percentiles, respectively. **e**, Percentage of SOX4^+^ and SOX4^−^ naïve CD4^+^ T cell subsets versus age of infants. Each line represents the best-fitted linear regression, with the shading showing the 95% confidence intervals. R represents the Pearson correlation coefficient. **f**, Percentage of SOX4^+^ and SOX4^−^ naïve CD8^+^ T cell subsets versus age of infants. **g**, Volcano plots showing differentially expressed genes between SOX4^+^ and SOX4^−^ naïve T cells for CD4^+^ (upper panel) and CD8^+^ lineages (lower panel). **h**, Boxplots showing log2CPM expression for selected differential genes in SOX4^+^ and SOX4^−^ naïve T cells for CD4^+^ (upper panels) and CD8^+^ lineages (lower panels). P-values were calculated using the Wilcoxon rank sum test (paired test). The upper and lower bounds represent the 75% and 25% percentiles, respectively.

26 genes were differentially expressed between SOX4^+^ and SOX4^−^ cells in CD4^+^ naïve T cells, whereas 36 were differentially expressed in CD8^+^ naïve T cells (**Extended Data Fig. 8g**). There was a significant overlap between the two lineages (**Extended Data Fig. 8h and Supplementary Table 6**). Genes upregulated in SOX4^+^ naïve T cells reflected a transcriptional signature associated with stemness and T cell development (*ID2*, *ID3, IKZF2*)^15,16^ (**Fig. 7g**). SOX4^+^ T cells also highly expressed *TCF7* and *LEF1* (**Fig. 7h**), key regulators of T cell development and maintenance. These TFs are components of the Wnt/β-catenin signaling pathway, which promotes stemness and self-renewal^17,18^. Pathway enrichment analysis revealed significant associations with thymocyte-related genes in both lineages (**Extended Data Fig. 8i**). SOX4^−^ naïve T cells, in contrast, highly expressed Th1-associated TF *STAT4* (**Supplementary Table 6**).

To investigate the epigenetic landscape of SOX4^+^ T cells across age groups, we performed single-nucleus ATAC-seq (snATAC-seq) on infants (n=8), children (n=4), young adults (n=5), and older adults (n=6). These data confirmed the robust separation of SOX4^+^ and SOX4^−^ subsets (**Fig. 8a**). As expected, chromatin accessibility around *SOX4* gene was significantly higher in SOX4^+^ naïve T cells compared to SOX4^−^ T cells (**Fig. 8b**). In alignment with the decline of SOX4^+^ cells with age, chromatin accessibility around *SOX4* gene promoter closed with age in both T cell lineages (**Fig. 8c**). Chromatin accessibility of genes linked to thymic emigration (*IKZF2*, *TOX*)^19,20^ and TGF-β signaling (*SMAD1*), an inducer of SOX4^21^ followed a similar trajectory, more accessible in infants and progressively closing with age (**Fig. 8c**). 1315 peaks were differentially accessible between SOX4^+^ and SOX4^−^ naïve T cells in CD4^+^ T cells and 1060 in CD8^+^ T cells (**Extended Data Fig. 9a and Supplementary Table 7**). SOX4^+^ CD4^+^ T cells exhibited increased chromatin accessibility in *CCR4* and *GATA3* (**Fig. 8e**), hallmark Th2 markers^22^, indicating an epigenetic Th2 differentiation bias in SOX4^+^ CD4^+^ T cells.

**Fig. 8.**
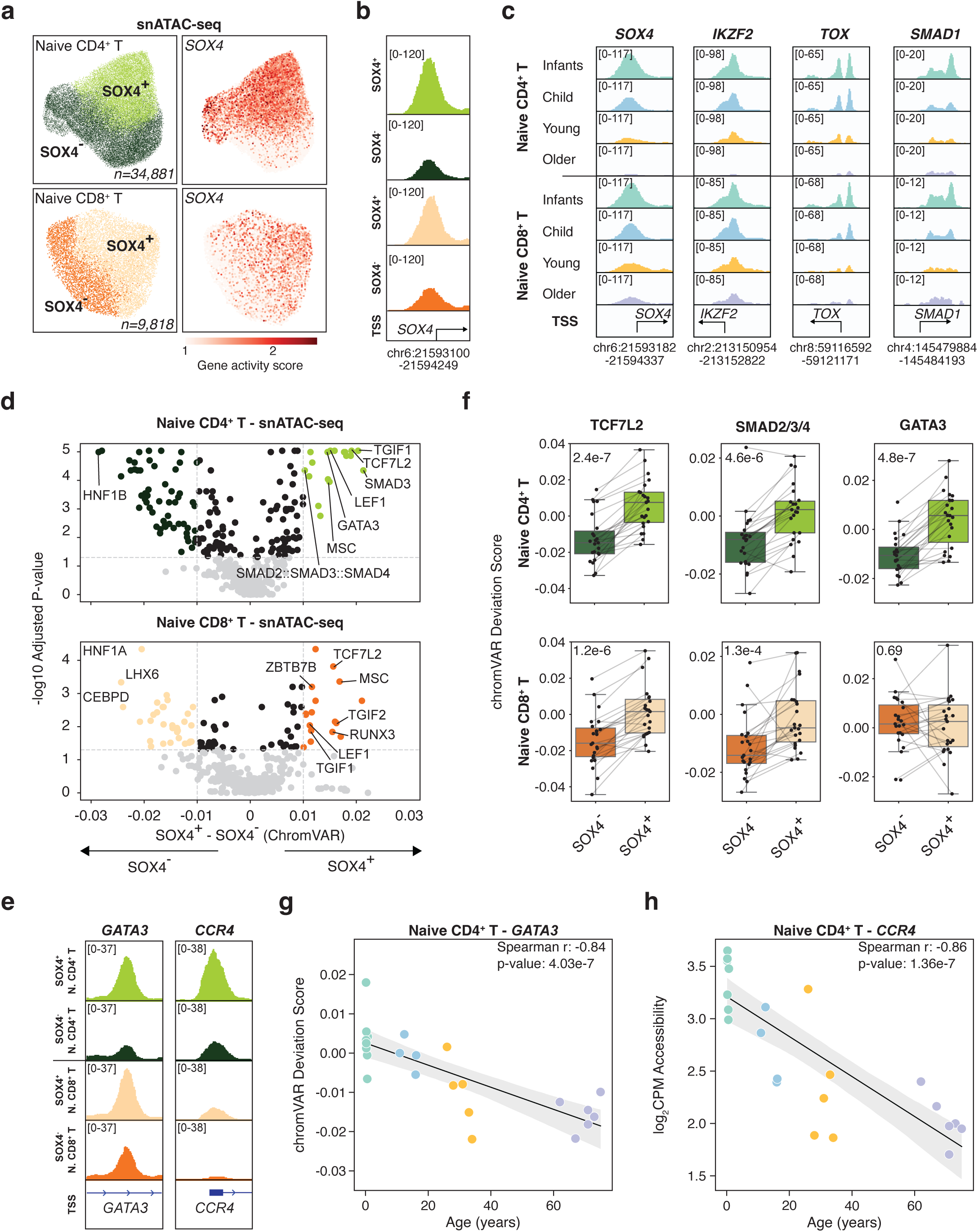
EpigenePcs landscape of SOX4^+^ T cells in infants. **a**, snATAC-seq data from 23 donors (8 infants, 4 children, 5 young and 6 older adults), UMAP plot representing SOX4^+^ and SOX4^−^ naïve T cell clusters in CD4^+^ (left panel) and CD8^+^ lineage (right panel) and SOX4 chromatin accessibility (gene activity score) for each subset. **b**, Genome browser to show accessibility around *SOX4* gene promoter for SOX4^+^ and SOX4^−^ naïve CD4^+^ and CD8^+^ T cells. **c**, Genome browser representing the normalized chromatin accessibility levels around the promoters of selected genes (*SOX4, IKZF2, TOX* and *SMAD1*) in total naïve CD4^+^ and CD8^+^ T cells across four age groups. Note the age-related decline in chromatin accessibility levels for these. **d**, Volcano plot showing differentially accessible peaks between SOX4^+^ and SOX4^−^ naïve T cells in CD4^+^ (upper panel) and CD8^+^ lineages (lower panel). **e**, Genome browser for CCR4 and GATA3 in SOX4^+^ and SOX4^−^ naïve CD4^+^ and CD8^+^ T cells. **f**, Box plots representing TCF7L2, SMAD2/3/4 and GATA3 chromVAR deviation scores in SOX4^+^ and SOX4^−^ in CD4^+^ and CD8^+^ T cells. **g**, Scatterplot representing chromVAR deviation scores for GATA3 in SOX4^+^ and SOX4^−^ naïve CD4^+^ T cells (y-axis) and age (x-axis). Each line represents the best-fitted linear regression, with the shading showing the 95% confidence intervals. R represents the Pearson correlation coefficient. **h**, Scatterplot representing log2 CPM accessibility for CCR4 in SOX4^+^ and SOX4^−^ naïve CD4^+^ T cells (y-axis) and age (x-axis). Each line represents the best-fitted linear regression, with the shading showing the 95% confidence intervals. R represents the Pearson correlation coefficient.

To further characterize the transcription factor (TF) landscape of SOX4^+^ naïve T cells, we performed ChromVAR analysis, which examines genome-wide chromatin accessibility at TF binding sites^23^. These analyses revealed that SOX4^+^ naïve CD4^+^ T cells exhibited preferential accessibility for TFs related to: (i) stemness (TCF7L, LEF1), (ii) TGF-b signaling (SMAD3, TGIF1), and (iii) Th2 commitment (GATA3, CCR4) (**Fig. 8d,e Extended Data Fig. 9d and Supplementary Table 8**). Additionally, SOX4^+^ naïve CD4^+^ T cells lacked accessibility for the Th1-associated TF TBX21 (T-Bet) (**Extended Data Fig. 9e**), which is in line with their Th2 epigenetic priming. SOX4^+^ naïve CD8^+^ T cells displayed a similar TF enrichment profile (TCF7L, LEF1, SMADs), but lacked CCR4 accessibility (**Fig. 8e,f)**. Binding site accessibility for TCF7L, SMADs, CCR4 and GATA3 declined with age (**Fig. 8g,h and Extended Data Fig. 9f**), consistent with the observed age-related decline in SOX4^+^ T cell frequencies.

To identify potential SOX4 binding targets, we selected peaks with the highest SOX4 motif score (>= 0.75 percentile) in naive CD4^+^ T and naive CD8^+^ T cells. A significant portion of DEGs between SOX4^+^ and SOX4^−^ cells (67-69%) contained a predicted SOX4 binding site (**Supplementary Table 9**), including SOX4 itself. This finding suggests that SOX4 may function as a key transcriptional regulator of stemness and SOX4^+^ T cell phenotype, including through direct autoregulation.

These data establish SOX4^+^ naïve T cells as a key feature of infant immunity, characterized by stemness and thymic emigrant signatures, and epigenetic priming toward Th2 differentiation.

## Discussion

We present a comprehensive single-cell map of peripheral blood mononuclear cells from 95 healthy individuals across the human lifespan. This dataset enabled us to delineate cross-sectional immune remodeling trajectories across age groups and revealed infancy as the most distinct period, characterized by unique immune features and significant remodeling in both innate and adaptive cell compartments. Major findings from this study included: (i) increased frequencies of monocytes and DCs with aging, (ii) more pronounced naïve-to-memory transitions in CD8⁺ than CD4⁺ T and B cells and (iii) an abundance of pDCs in infant DCs, (iv) the presence of constitutive ISG^hi^ T and B cells in early life, and (v) high frequencies of SOX4^+^ naïve CD4^+^ and CD8^+^ T cells in infants.

All myeloid subsets including monocytes and DCs, increased with age, with the most pronounced expansion occurring during the transition to childhood. However, cDC1, which constitute ∼10% of DC subsets and are responsible for presenting antigenic peptides to CD8^+^ T cells^24,25^, displayed a distinct pattern by expanding later, in adulthood, unlike other DC subsets (*e.g.,* cDC2 and moDC). This delayed expansion may be linked to age-associated decline in naïve CD8^+^ T cells and the concurrent rise in cytotoxic CD8^+^ T cells.

Within lymphocytes, as expected, we observed a transition from naïve to memory phenotypes with age. This shift was more pronounced in the CD8⁺ T cell compartment than in CD4⁺ T cells, which might stem from three possible mechanisms: (i) cumulative effects of persistent viral infections (*e.g.,* CMV, EBV, VZV)^26–28^, (ii) age-related differences in thymic output favoring CD4⁺ T cells^29,30^, and (iii) differences in activation and maintenance programs between CD4⁺ and CD8⁺ T cells^31^. Among CD8⁺ T cell subsets, TEMRA and GZMK⁺ populations showed the greatest expansion, constituting ∼14% of PBMCs in older adults, which is in line with previous reports^7^. In contrast, CD4⁺ memory T cell expansion was more modest, and we did not observe significant age-associated shifts within HLA-DR⁺ or Th2 cells^8^.

MAIT and γδ T cell frequencies increased in children and young adults but declined in older adults, following a characteristic ‘rise and fall’ pattern that has been documented in other cohorts^32,33^. This pattern might be associated to age-dependent microbial exposures, as MAIT cells respond to microbial-derived metabolites, while γδ T cells are activated by phosphoantigens produced by microbes^34^. Their expansion during childhood may reflect increased microbial priming during this window^35^, while the decline with age could be due to reduced microbial diversity and/or compromised barrier integrity^34^. Hormonal changes during adolescence and menopause/andropause, which affect commensal communities^36^, may also contribute to these trends.

Strikingly, several features of infant immunity were associated with antiviral responses. In infants, plasmacytoid dendritic cells (pDCs) comprised ∼50% of the DC pool, compared to ∼28% in adults. Given their potent type I interferon production, abundance of pDCs may highlight prompt innate responses to compensate for the still-developing adaptive immunity^1,37^.

Additionally, a greater proportion of infant T and B cells constitutively expressed interferon-stimulated genes (ISGs) even in the absence of known acute infection or recent vaccination (**Extended Data Fig. 10**). This ISG-high state may reflect a pre-activated anti-viral program^38,39,40,13^, potentially compensating for the immaturity and limited repertoire of the infant’s adaptive immune system.

Infant PBMC had high frequency of SOX4⁺ naïve T cells in both the CD4⁺ and CD8⁺ compartments—accounting for ∼30% of infant naïve T cells, a population that sharply declined with development and aging. SOX4 expression has been linked to recent thymic emigrants (RTEs)^20^, which may possess distinct effector functions, including a reduced capacity for Th1 polarization^41^ and a bias towards regulatory T cell generation^42^. Our epigenetic profiling of SOX4^+^ T cells revealed increased chromatin accessibility at the *GATA3* and *CCR4* loci, suggesting an epigenetic Th2 bias, which may contribute to the predisposition of infants toward Th2 responses and increased susceptibility to allergy upon early-life allergen exposure^42^. Indeed, recent studies showed a Th2 skew in infant CD4^+^ T cell responses in both humans^43,44^ and mice^45^. Other studies reported that SOX4⁺ naïve T cells are functionally hyporesponsive to activation *in vitro*^20,46^, however, their *in vivo* roles and responses warrant further investigation. A previous report showed that SOX4 promote the formation of CXCL13-producing CD4^+^ helper T cells at sites of local inflammation^47^. This suggests that a subset of infant SOX4^+^ CD4^+^ T cells may contribute to nascent humoral immunity by promoting CXCL13-producing helper T cell development.

SOX4 expression was not restricted to conventional naïve T cells, as a SOX4⁺ γδ T cell population with naïve-like features was abundant in infants and declined with age. A SOX4^+^ memory CD4^+^ T cell subset was also identified, resembling previously described tissue-resident cells-expressing *LEF1* and *IKZF2* (Helios) that are detected in multiple tissues in children (0-10 years)^48^. These cells may represent a circulating reservoir of stem-like memory T cells involved in establishing and maintaining tissue niches^48^.

Together, our findings underscore the dynamics and unique features of infant immunity, including enhanced antiviral preparedness and the abundance of SOX4^+^ naïve T cells, while also providing a comprehensive map of immune remodeling across the human lifespan. This resource lays a foundation for better understanding age-specific immune vulnerabilities and could inform strategies to enhance immune function in both early and late life.

### Limitations of the study

While our study provides a comprehensive single-cell map of immune changes across the human lifespan, several limitations should be acknowledged. Our analysis is limited to PBMCs, which do not capture tissue-resident immune cells. Future studies integrating tissue-specific immune profiles could provide a more holistic understanding of immune development and aging. Our cross-sectional design limits the ability to directly infer longitudinal immune dynamics, and prospective longitudinal studies within age groups are needed to confirm age-related trends and transitions. On the other hand, our strategy including individuals across well characterized age groups allowed us to identify major differences in cell population frequencies. Indeed, the age-associated shifts in various immune subsets reported in this study call for further mechanistic investigations, particularly exploring their connections with changes in metabolites and microbiome compositions.

Despite these limitations, our study included a unique cohort of healthy individuals spanning from 2 months to 88 years of age and offers valuable insights into the cellular and molecular changes that shape the human immune system across the lifespan highlighting unique features of the infant immune system.

## Methods

### Study design

#### 1 – Pediatric cohorts

Healthy infants were enrolled at primary care offices during well-child visits or in the operating room while undergoing minor scheduled surgical procedures not involving the respiratory tract, as previously described^13,40,44^. Written informed consent was obtained from all children’s guardians before study participation. The study was approved by the Nationwide Children’s Hospital (NCH) IRB (18-00591).

#### 2 – Adult cohorts

Healthy young adults were recruited during 2017-2018 from communities surrounding Hartford, Connecticut, USA. Healthy older adults were recruited during 2017-2018 from communities surrounding Hartford, Connecticut, USA, and Sudbury, Ontario, Canada. Both young and older adults were free from confounding autoimmune diseases, were not undergoing treatments that might influence immune function and composition (*e.g.,* immunotherapy) and were not recently vaccinated prior to blood collection, as previously described^49,50^. The study protocols were approved by the Institutional Review Board of the University of Connecticut Health Center (UConn Health IRB #20-130-J-1, #16-071J or JAX IRB #2019-006-JGM) and the Health Sciences North Research Ethics Board (#985). Written informed consent was obtained from all participants prior to inclusion in the study.

Detailed information on participant demographics is provided in Supplementary Table 1.

### Blood preparation for single cell RNA sequencing (scRNA-seq)

PBMCs were thawed quickly at 37°C and into DMEM supplemented with 10% FBS. Cells were quickly spun down at 400 g, for 10 min. Cells were washed once with 1 x PBS supplemented with 0.04% BSA and finally re-suspended in 1xPBS with 0.04% BSA. Viability was determined using trypan blue staining and measured on a Countess FLII. Briefly, 12,000 cells were loaded for capture onto the Chromium System using the v2 single cell reagent kit (10X Genomics). Following capture and lysis, cDNA was synthesized and amplified (12 cycles) as per manufacturer’s protocol (10X Genomics). The amplified cDNA was used to construct an Illumina sequencing library and sequenced on a single lane of a HiSeq 4000.

### Blood preparation for for snATAC-seq

For single nucleus ATAC sequencing (snATAC-seq) experiments, viable single cell suspensions from each sample were used to generate snATAC-seq data using the 10x Chromium platform according to the manufacturer’s protocols (10x Genomics, protocols CG000169; CG000168). Briefly, >100,000 cells from each sample were centrifuged and the supernatant was removed without disrupting the cell pellet. Lysis Buffer was added for 5 minutes on ice to generate isolated and permeabilized nuclei, and the lysis reaction was quenched by dilution with Wash Buffer. After centrifugation to collect the washed nuclei, diluted Nuclei Buffer was used to re-suspend nuclei at the desired nuclei concentration as determined using a Countess II FL Automated Cell Counter and combined with ATAC Buffer and ATAC Enzyme to form a Transposition Mix. Transposed nuclei were immediately combined with Barcoding Reagent, Reducing Agent B and Barcoding Enzyme and loaded onto a 10x Chromium Chip H for droplet generation followed by library construction. The barcoded sequencing libraries were subjected to bead clean-up and checked for quality on an Agilent 4200 TapeStation, quantified by KAPA qPCR, and pooled for sequencing on an Illumina NovaSeq 6000 (2×50bp libraries).

### Single-cell Raw data processing and data integration

Illumina basecall files (*.bcl) were converted to fastq files using cellranger v3.0.2, which uses bcl2fastq v2.17.1.14. FASTQ files were then aligned to hg19 genome and transcriptome using the cellranger v3.0.2 pipeline, which generates a gene – cell expression matrix. The samples were combined using *cellranger aggr* from cellranger, which aggregates outputs from multiple runs, normalizing them to the same sequencing depth (*normalize=mapped*) and then re-computing the gene-barcode matrices and analysis on the combined data. The sequencing information for each sample included in the study (*e.g*., number of reads per cell or UMI counts per cell) are shown in **Supplementary Table 2.**

### Scrublet for multiplet prediction and removal

Generally, we expected about %2-8 of the cells to be hybrid transcriptomes or multiplets, occurring when two or more cells are captured within the same microfluidic droplet and are tagged with the same barcode. Such artificial multiplets can confound downstream analyses. We applied Scrublet^51^ python package to remove the putative multiplets. Scrublet assigns each measured transcriptome a ‘multiplet score’, which indicates the probability of being a hybrid transcriptome. Multiplet scores were determined for each sample (using the raw data). Before/after multiplet removal, the number of cells was 715,910/687,819. After visual inspection the number of cells was 581,724.

### Single-cell data preprocessing, clustering, and annotation

The cleaned (after multiplet removal using scrublet^51^ aggregated matrices were fed into the Python-based Scanpy^52^ workflow (https://scanpy.readthedocs.io/en/stable/), which includes preprocessing, visualization, clustering and differential expression testing.

### Quality control and cell filtering

We applied the following filtering parameters: (i) all genes that were not detected in ≥ 3 cells were discarded, using *pp.filter_genes* function, (ii) cells with less than 400 total unique transcripts were removed prior to downstream analysis using *pp.filter_cells* function, (iii) cells in which > 25% of the transcripts mapped to the mitochondrial genes were filtered out, as this can be a marker of poor-quality cells and (iv) cells with > 3,500 genes were considered outliers and discarded.

### Data normalization, log transformation and scaling

We normalized the data using the *pp.normalize_per_cell* function. Thus, library-size normalization was performed based on the gene expression for each barcode by scaling the total number of reads per cell to 10,000. We log-transformed the data using the *pp.log1p* function and scaled to unit variance using *pp.scale* function (with the following parameters: max_value=10). The 1,173 highly variable genes (HVG) were identified using *filter_genes_dispersion* function (with the following parameters: min_mean=0.0125, max_mean=3, min_disp=0.5) (**Extended Data Fig. 1c**).

### Linear dimensional reduction using PCA

To reduce the dimensionality of the data, we ran principal component analysis (PCA) using *tl.pca* function, which reveals the main axes of variation and denoises the data. The contribution of each PCs to the total variance was assessed using *pl.pca_variance_ratio* function.

### Neighborhood graph computing, embedding and clustering

The neighborhood graph of cells was computed based on the PCA representation of the data matrix, using *pp.neighbors* function (with the following parameters: n_neighbors=10, n_pcs=40). The neighborhood graph was then embedded using UMAP^53^ (*tl.umap* function) and visualized using *pl.umap* function. We finally used the Leiden graph-based clustering using *tl.leiden* function with resolution=1.2 (stored in ‘Res1_2_BC’ column).

### Batch effect correction

To account for technical source of variation due to chemistry (V2 *vs.* V3) and 10X runs (eight samples per run), we applied Harmony^54^ for batch correction (**Extended Data Fig. 1d,e**), as detailed in the following commands:

*pca = Anndata_object.obsm[’X_pca’]*

*batch = Anndata_object.obs*

*hem <-HarmonyMatrix(pca, batch,c(’Chemistry_10x’,’runs_10x’), do_pca=FALSE)*

*hem = data.frame(hem)*

*har_corrected= Anndata_object.copy()*

*har*_corrected_object.*obsm[’X_pca’] = hem.values*

All the analyses presented in this study were perfomed on the corrected data. UMAP visualization and Leiden clustering were then computed on the corrected data using the following parameters:

*sc.tl.umap(har_corrected_object, min_dist=0.3, n_components=3) sc.tl.leiden(har_corrected_object, resolution=1.2, key_added= ‘Res1_2_Harm’)*

### Finding marker genes/evaluation of cluster identity

To annotate the clusters generated from the BBKNN corrected object, we used both differential expression analysis between clusters and classification based on putative marker gene expression. We applied *tl.rank_genes_groups* function to compute a ranking for the differential genes in each subcluster, comparing each cluster to the rest of the cell using Wilcoxon test. We only considerate subclusters that showed distinct transcriptomic programs. The top 100 marker genes for cluster and subclusters are included in **Supplementary Table. 3 & 4**. The top 10 marker genes were visualized using the *sc.pl.rank_genes_groups_matrixplot* function.

### Subclustering parameters and data cleaning

We only considered SC defined by distinct gene sets, by merging similar ones. Based on the number of cells and to avoid over-clustering, we used different clustering resolutions. Subcluser cell frequencies across donors are available in **Supplementary Table 5**. A script showing the subclustering process can be found here: https://github.com/dnehar/Lifespan_project.

### Differential gene expression analysis for SOX4^+^ and SOX4^−^ comparisons

Aggregated raw gene expression count matrices were obtained for each annotated subset. For each comparison, samples with fewer than 50 cells in the respective population were excluded. Genes were excluded if fewer than 33% of samples had expression (> 0) in more than 10% of cells for both groups in the comparison. Sample and gene filtered count matrices were then normalized using *filterByExpr* and *calcNormFactors* functions in EdgeR^55^ (version 4.1.26). Differential expression analyses were performed using *estimateDisp* and *glmQLFTest* functions, blocking on sample identifier to perform pairwise comparisons within the same donor. P-values were adjusted using p.adjust (method=fdr) and differentially expressed genes were identified using abs(logFC) > 0.5 and adjusted p-value < 0.05.

### snATAC-seq data processing

snATAC-seq reads were aligned (10x Genomics GRCh38 reference 2020-A) and processed using Cell Ranger ATAC v2.1.0. High quality single nucleus data was selected based on the following criteria: percent of reads within exclusion list regions < 5%, nucleosome signal < 4, percent of fragments in peaks > 15%, number of peak region fragments > 3000, percent fragments at transcription start sites > 10%, and percent mitochondrial read fragments < 10%. Multiplets were identified by applying AMULET^56^ on each sample independently using FDR < 5%. Multiplet scores were used to identify and remove clusters with high frequency multiplet frequencies. Subsequently, all multiplets identified by AMULET were excluded after first pass filtering to account for homotypic multiplets.

To define a unified set of peaks (genomic regions) for clustering, individual sample peaks from Cell Ranger ATAC were combined by merging peaks if they overlapped by 1 base pair. Peaks with lengths < 20bp; > 10,000bp, falling on chromosome Y, or represented by fewer than 3 samples were excluded from further analysis. Read count matrices were generated from the remaining peaks for each sample. Read count matrices were concatenated and normalized by applying a python reimplementation of the term frequency inverse document frequency (TF-IDF) normalization method in Signac^57^. Scanpy objects were generated using this matrix, and singular value decomposition (SVD) of the combined TF-IDF normalized matrix was performed using the *pca* function in Scanpy with zero_center=False to perform truncated SVD. To measure read counts at transcription start sites (TSS) and gene bodies, gene activity scores were computed using a python reimplementation of gene activity score calculations in Signac. TSS and transcription termination sites from UCSC hg38 Refflat database^58^ were used as gene references. Quantifications included reads within the gene body and 2000 base pairs upstream from the transcription start sites.

### snATAC-seq label transfer from scRNA-seq for SOX4^+^ naïve T subsets

Nuclei/cells from snATAC-seq and scRNA-seq were divided into CD4^+^ T and CD8^+^ T subset based on gene activity scores and gene expression values respectively. Within each subset, cluster annotations from scRNA-seq were transferred onto snATAC-seq cells using *FindTransferAnchors* with reduction=’CCA’ and *TransferData* functions in Signac (version 1.7.0)^57^ and Seurat (version 4.1.1)^59^. Nuclei were then assigned scRNA-seq annotations based on highest probability scores. Based on these annotations, clusters in snATAC-seq data were assigned to reflect most cell predictions within the cluster.

### Differential accessibility analysis for SOX4^+^ and SOX4^−^ comparisons

Aggregated raw read count matrices were obtained for each annotated subset using peaks called for the respective subset to define the genomic regions used in these matrices. For each comparison, samples with fewer than 50 cells in the respective population were excluded. Sample and gene filtered count matrices were then normalized using *filterByExpr* and *calcNormFactors* functions in EdgeR^55^ (version 4.1.26). Differential accessibility analyses were performed using *estimateDisp* and *glmQLFTest* functions blocking on sample identifier to perform pairwise comparisons within the same donor. P-values were adjusted using p.adjust (method=fdr) and differentially expressed genes were identified using abs(logFC) > 0.5 and adjusted p-value < 0.05.

### Transcription factor motif binding site accessibility for SOX4^+^ and SOX4^−^ comparisons

Transcription factor motif binding sites accessibility deviations were calculated using chromVAR version 1.16.0^23^. For each PBMC subset, genomic regions for chromVAR analysis were selected by extending MACS2 summit locations ± 250 base pairs (bp). Read count matrices were then generated based on single nucleus read counts within these regions. TF motif deviations were then calculated using *computeDeviations* in chromVAR, using motifs from *getJasparMotifs()*, (CORE set JASPAR 2016^60^, homo sapiens). Motifs were selected based on TFs with expression > 0 within the respective PBMC subset. Paired tests were applied on median motif deviation scores within the same donor (SOX4^+^ and SOX4^−^ cells) using Wilcoxon signed rank test. P-values were adjusted for multiple comparisons within each PBMC subset using Benjamini/Hochberg procedure. Finally, significant motifs were selected using adjusted p-values < 0.05.

### SOX4 target peak identification

SOX4 motif scores were obtained using the *annotatePeaks* function in HOMER (v4.11.1)^61^ using the –mscore option and the SOX4 position weight matrix from Jaspar2020 database. Peaks within the 75^th^ percentile of the top motif log odds score were selected as SOX4 target peaks for downstream analysis.

### Statistical analysis

Statistical analysis was performed using R/4.0.2. Tests were used to determine data distribution and depending on the normality of the data, comparisons were performed using the Student t test (for two groups, parametric) or the non-parametric the Wilcoxon signed rank test (for two groups, paired) with two-tailed P values unless otherwise stated. Differences were considered when P<0.05 (*), P < 0.01 (**), P < 0.001 (***) and P < 0.0001 (****).

### Python module versions

scanpy==1.7.1 anndata==0.7.5 umap==0.4.6 numpy==1.19.2 scipy==1.5.2 pandas==1.1.3 scikit-learn==0.24.1 statsmodels==0.12.2 python-igraph==0.8.3 leidenalg==0.8.3

## Data availability

The processed scRNA-seq data generated in this study have been deposited in the Gene Expression Omnibus (GEO) database under accession code: GSE233321. The fastq files have been deposited in the dbGAP database under accession code: phs003259.v1.p1 (will be released upon acceptance). Sorted γδ T cell scRNA-seq data from newborns and adult can be found here: GSE149356.

## Code availability

Scripts used to process and analyze the data are available on GitHub: https://github.com/dnehar/Lifespan_project. We provide open access to our data via an interactive web app R Shiny app: https://dnehar.shinyapps.io/LS_app/

## Acknowledgments

We thank M. Collet and O. Bart for help with dbGAP data upload, T. Helenius for help in scientific writing and research staff in the UConn Center on Aging for their help in recruitment and sample collection and the JAX Genomic Technologies and Single Cell cores for help with generating the sequencing data. The JAX single-cell service is supported, in part, by the JAX Cancer Center P30 CA034196 (to K.P., D.U. and J.F.B). We thank members of the Ucar lab for critical feedback during the progress of the study. This study was made possible by generous financial support from NIH grants under award numbers U01 AI 131386 (to O.R. and J.F.B), R01AG052608 (to J.F.B), U01AI165452 (to D.U. and G.A.K.), P30AG067988 Older Americans Independence Pepper Center (to G.A.K.) and R01AI142086 (to D.U. and J.F.B). U19 AI168632 (to VP, DU, AM and OR). Opinions, interpretations, conclusions and recommendations are solely the responsibility of the authors and do not necessarily represent the official views of the NIH.

## Author contributions

D.N-B. designed the experiments, led the data analyses, interpretation, and figure generation. A.T. led the ATAC-seq data analysis, helped with data analyses and data interpretation. A.E. helped with data analyses and data interpretation. R.M. processed the PBMCs and contributed to data generation. G.E, J.G and U.B helped interpreting the data. C.P.V., V.P., G.A.K., O.R and A.M. coordinated the clinical sample collection and interpreted the data. D.U, O.R and J.F.B. conceived, supervised the study and interpreted the data. D.N-B. and D.U. wrote the first draft of the manuscript. All authors reviewed and contributed to the final manuscript.

## Ethics declarations

In the early stage of the study, J.F.B. was a member of the BOD and SAB of Neovacs and Ascend Biopharma, and a SAB member of Cue Biopharma. Now, JFB is the Founder of Immunoledge LLC, an entity designed to advise biotech start-ups. In this role, JFB serves as CIO of Georgiamune, SAB member of Metis Therapeutics, and Adviser to the JAX. O.R. has received research grants from the NIH, Gates Foundation and Merck; and fees for participation in advisory boards and educational lectures from Merck, Sanofi-Pasteur, Moderna and Pfizer.

The remaining authors declare no competing interests.

**Extended Data Fig. 1.**
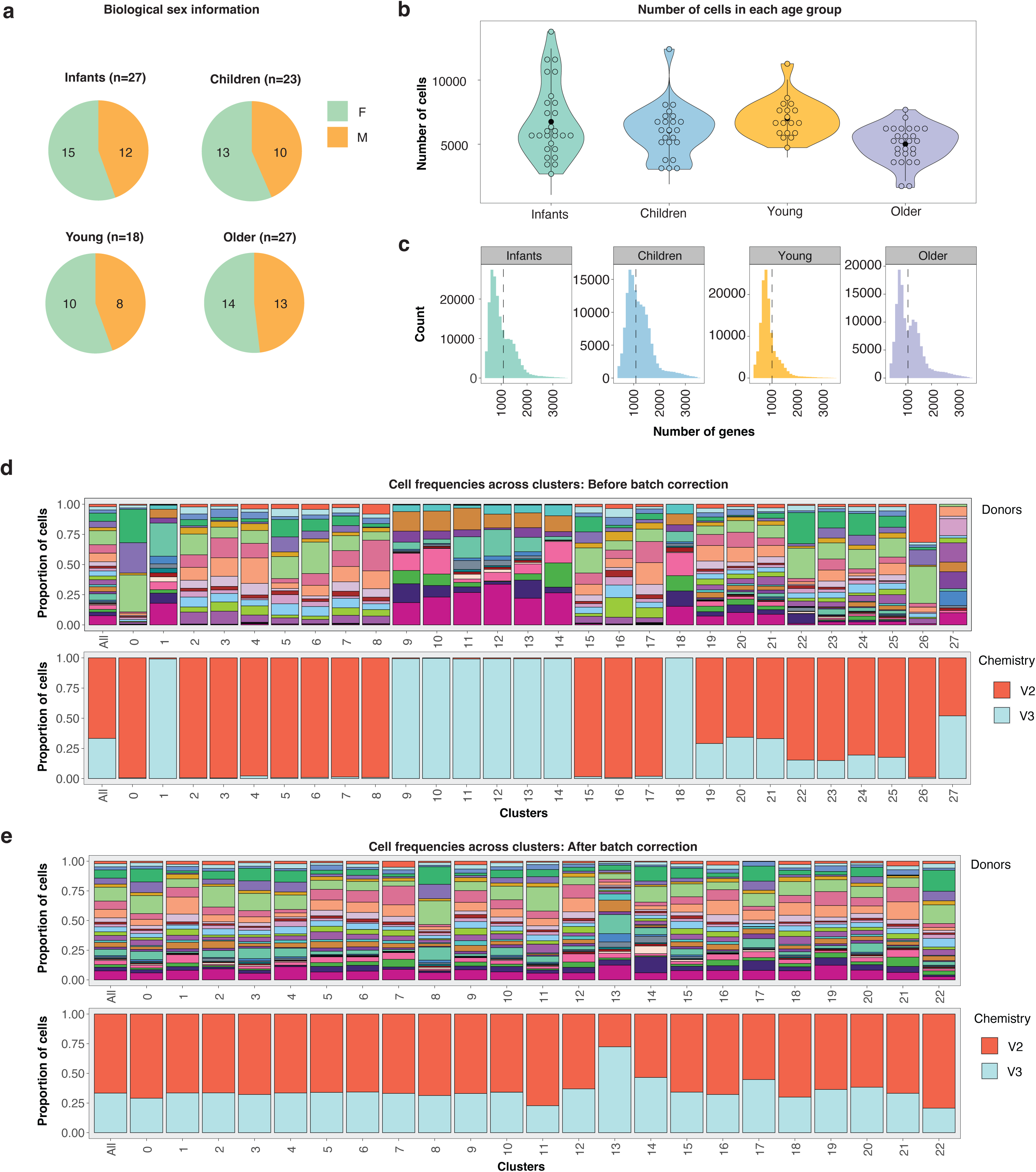
Quality control information. **a**, Biological sex repartition in each age group. **b**, Violin plots representing the number of cells across age groups. **c**, Number of detected genes per cell across age groups. **d-e**, Bar plot representing cell compositions contributed by donors (upper panel) and V2/V3 10X chemistry (lower panel), before (**d**) and after (**e**) batch effect correction.

**Extended Data Fig. 2.**
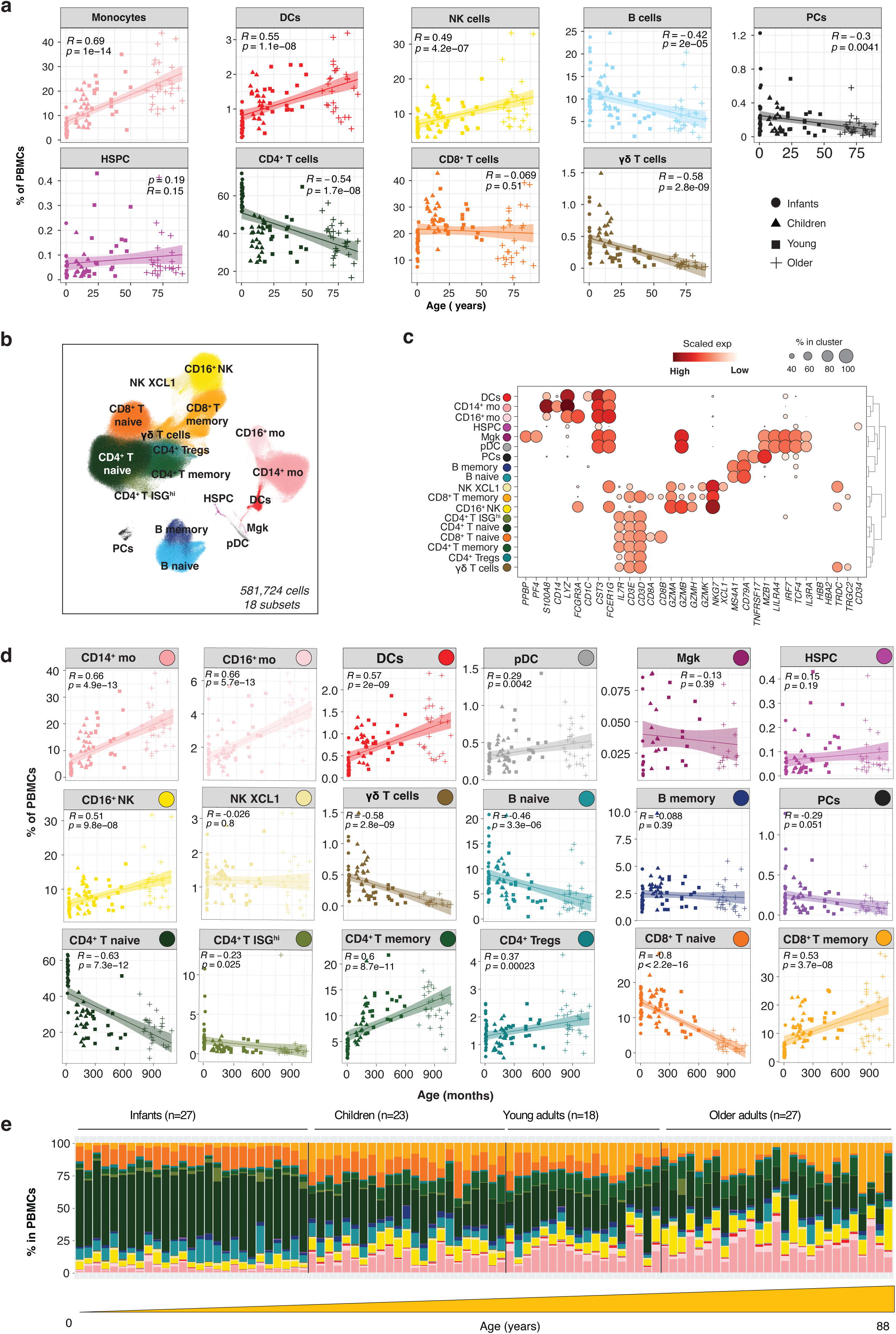
Low resolution clustering. **a**, Scatterplot representing the frequencies of immune subsets (n=9) versus age. Each line represents the best-fitted linear regression, with the shading showing the 95% confidence intervals. R represents the Pearson correlation coefficient. **b**, UMAP plot representing 581,724 PBMCs colored by immune subsets (n=18). **c**, Dot plot represents expression values of selected genes (x-axis) across each cluster (y-axis). Dot size represents the percentage of cells expressing the marker of interest. Color intensity indicates the mean expression within expressing cells. **d**, Scatterplot representing the frequencies of immune subsets (n=18) versus age. Each line represents the best-fitted linear regression, with the shading showing the 95% confidence intervals. R represents the Pearson correlation coefficient. **e**, Barplot showing the frequencies of each immune subsets (n=18) across individuals (n=95).

**Extended Data Fig. 3.**
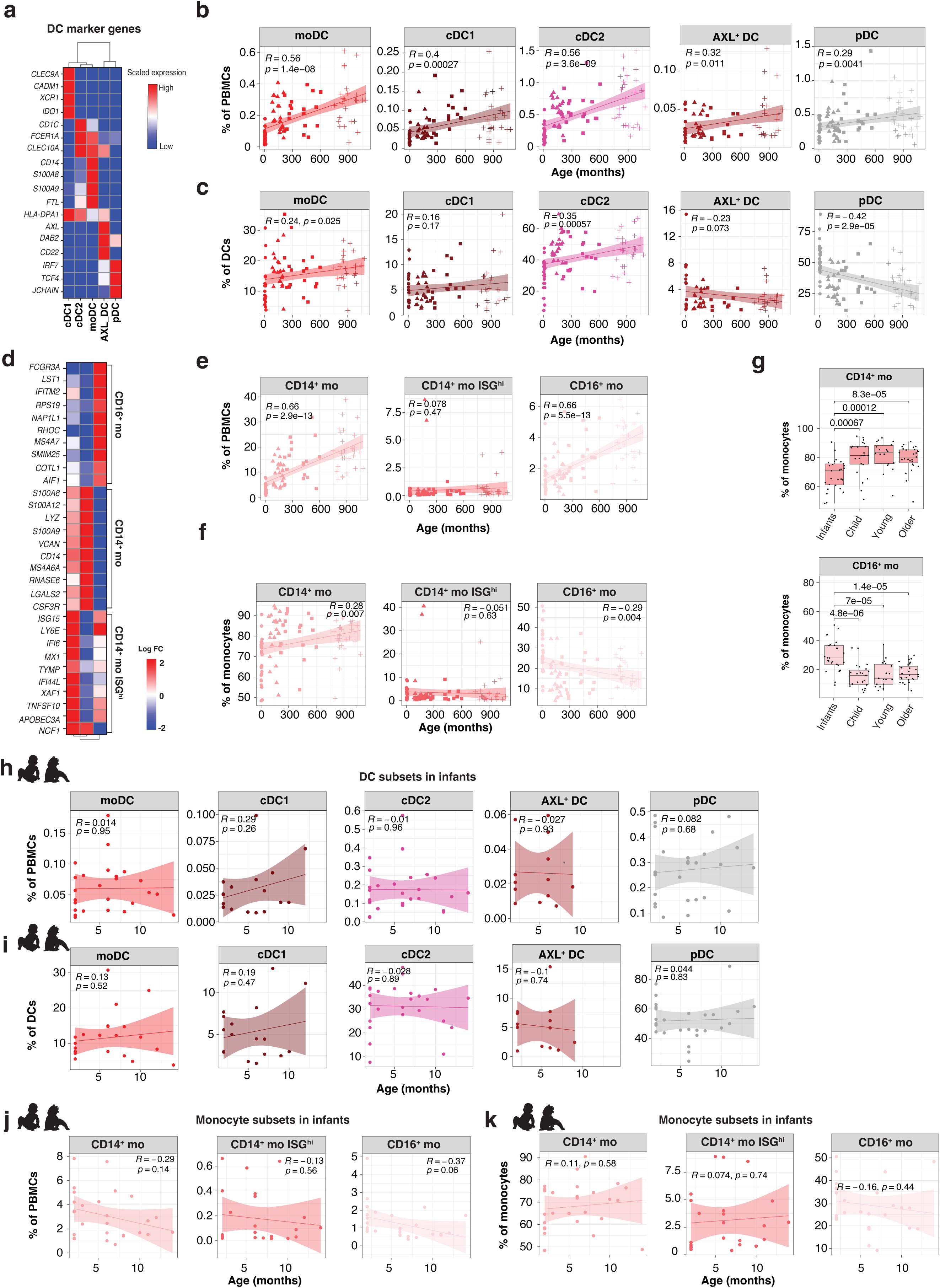
Myeloid cells – Related to Fig. 2. **a**, Heat map representing log fold change values of selected marker genes defining each of the DC subsets**. b**, Scatterplot representing the frequencies of each DC subset in PBMCs (y-axis) and age (x-axis). Each line represents the best-fitted linear regression, with the shading showing the 95% confidence intervals. R represents the Pearson correlation coefficient. **c**, Scatterplot representing the percentage of each DC subset in lineage (y-axis) and age (x-axis). Each line represents the best-fitted linear regression, with the shading showing the 95% confidence intervals. R represents the Pearson correlation coefficient. **d**, Heat map representing log fold change values of selected marker genes defining each of the monocyte subsets. **e**, Scatterplot representing the percentage of each monocyte subset in PBMCs (y-axis) and age (x-axis). Each line represents the best-fitted linear regression, with the shading showing the 95% confidence intervals. R represents the Pearson correlation coefficient. **f**, Scatterplot representing the percentage of each monocyte subset in lineage (y-axis) and age (x-axis). Each line represents the best-fitted linear regression, with the shading showing the 95% confidence intervals. R represents the Pearson correlation coefficient. **g**, Boxplots showing distribution of CD14^+^ and CD16^+^ monocytes within monocytes across age groups. **h**, as in (**b**) in infants only. **i**, as in (**c**) in infants only. **j**, as in (**e**) in infants only. **k**, as in (**f**) in infants only.

**Extended Data Fig. 4.**
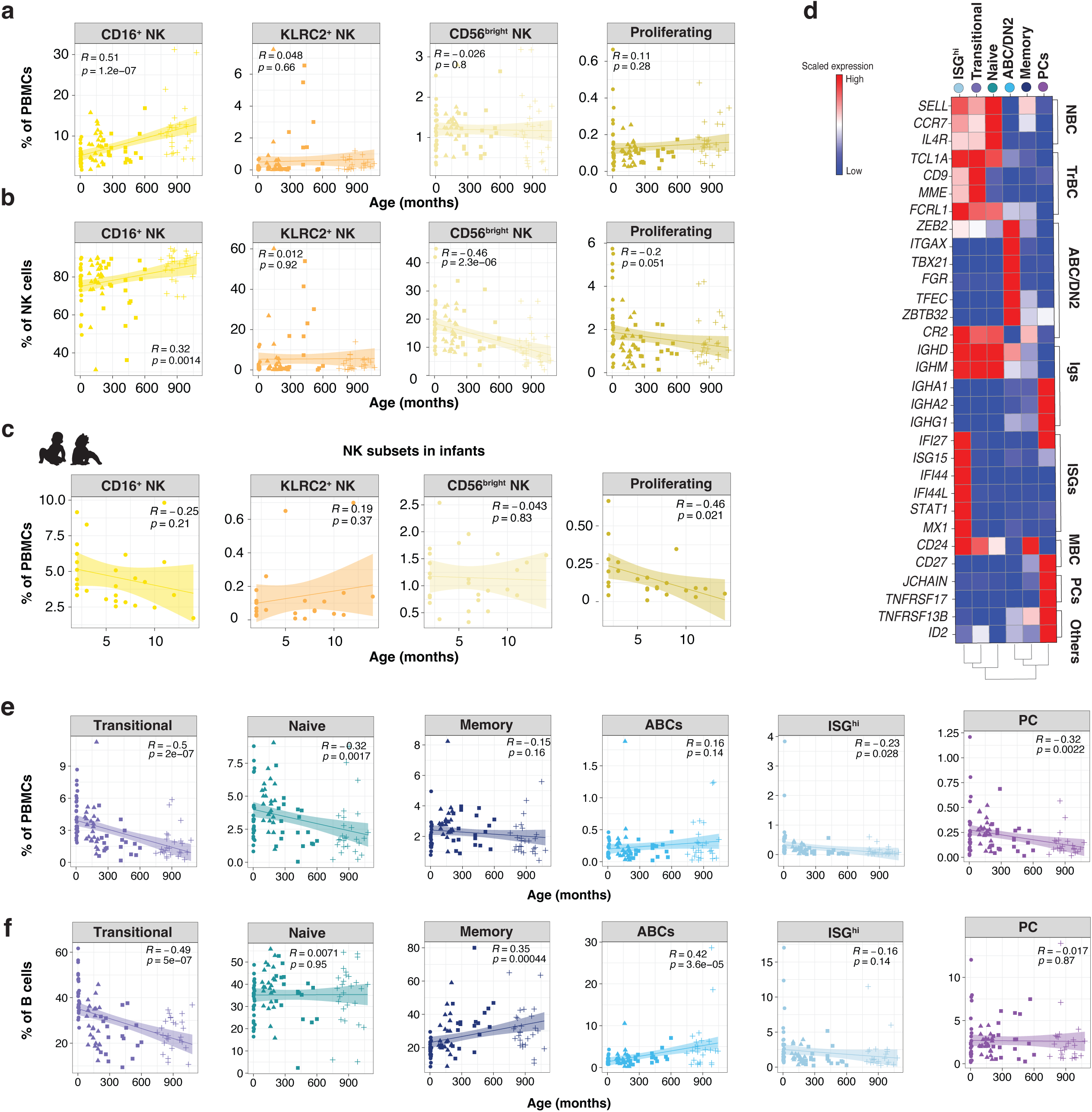
NK and B cells – Related to Fig. 3 & 4. **a**, Scatterplot representing the frequencies of each NK subset in PBMCs (y-axis) and age (x-axis). Each line represents the best-fitted linear regression, with the shading showing the 95% confidence intervals. R represents the Pearson correlation coefficient. **b**, Scatterplot representing the frequencies of each NK subset in lineage (y-axis) and age (x-axis). Each line represents the best-fitted linear regression, with the shading showing the 95% confidence intervals. R represents the Pearson correlation coefficient. **c**, as in (**a**) in infants only. **d**, Heat map representing log fold change values of selected marker genes defining each of the B cell subsets. **e**, As in **(a)** in B cells. **f**, As in **(b)** in B cells.

**Extended Data Fig. 5.**
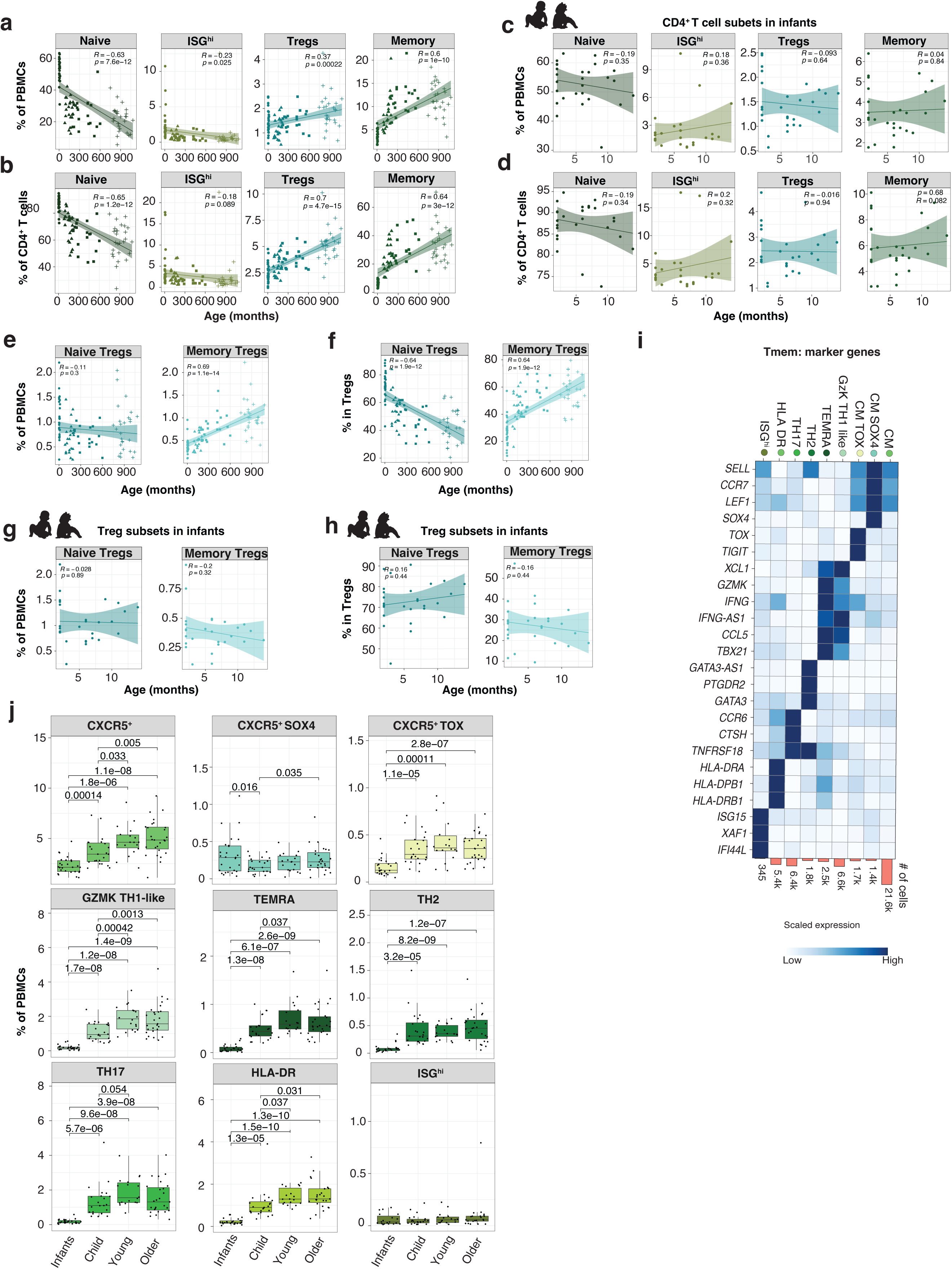
CD4^+^ T cells – Related to Fig. 5. **a**, Scatterplot representing the frequencies of in each CD4^+^ T cell subsets in PBMCs (y-axis) and age (x-axis). Each line represents the best-fitted linear regression, with the shading showing the 95% confidence intervals. R represents the Pearson correlation coefficient. **b**, Scatterplot representing the frequencies of in each CD4^+^ T cell subsets in total CD4^+^ T cells (y-axis) and age (x-axis). Each line represents the best-fitted linear regression, with the shading showing the 95% confidence intervals. R represents the Pearson correlation coefficient. **c**, As in **(a)** in infant only. **d**, As in **(b)** in infant only. **e**, Scatterplot representing the percentage of Treg subsets in PBMCs (y-axis) and age (x-axis). Each line represents the best-fitted linear regression, with the shading showing the 95% confidence intervals. R represents the Pearson correlation coefficient. **f**, Scatterplot representing the percentage of Treg subsets in total Tregs (y-axis) and age (x-axis). Each line represents the best-fitted linear regression, with the shading showing the 95% confidence intervals. R represents the Pearson correlation coefficient. **g**, As in **(e)** in infant only. **h**, As in **(f)** in infant only. **i**, Heat map representing scaled expression values of selected marker genes defining each of the Tmem subsets. **j**, Boxplots showing distribution of Tmem subsets within PBMCs across age groups.

**Extended Data Fig. 6.**
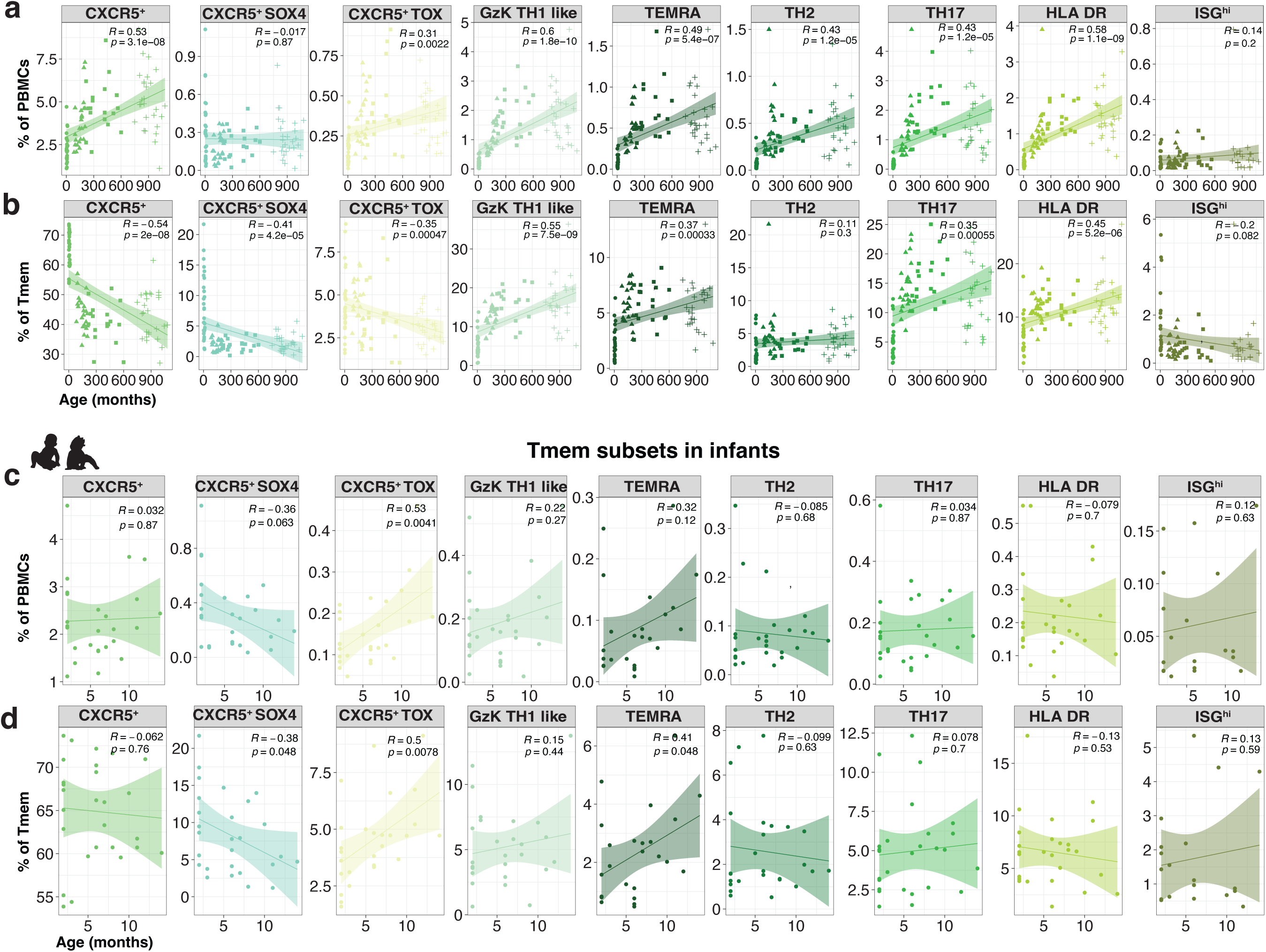
CD4^+^ T cells – Related to Fig. 5. **a**, Scatterplot representing the frequencies of each Tmem subsets in PBMCs (y-axis) and age (x-axis). Each line represents the best-fitted linear regression, with the shading showing the 95% confidence intervals. R represents the Pearson correlation coefficient. **b**, Scatterplot representing the frequencies of in each Tmem subset in lineage (y-axis) and age (x-axis). Each line represents the best-fitted linear regression, with the shading showing the 95% confidence intervals. R represents the Pearson correlation coefficient. **c**, As in **(a)** in infant only. **d**, As in **(b)** in infant only.

**Extended Data Fig. 7.**
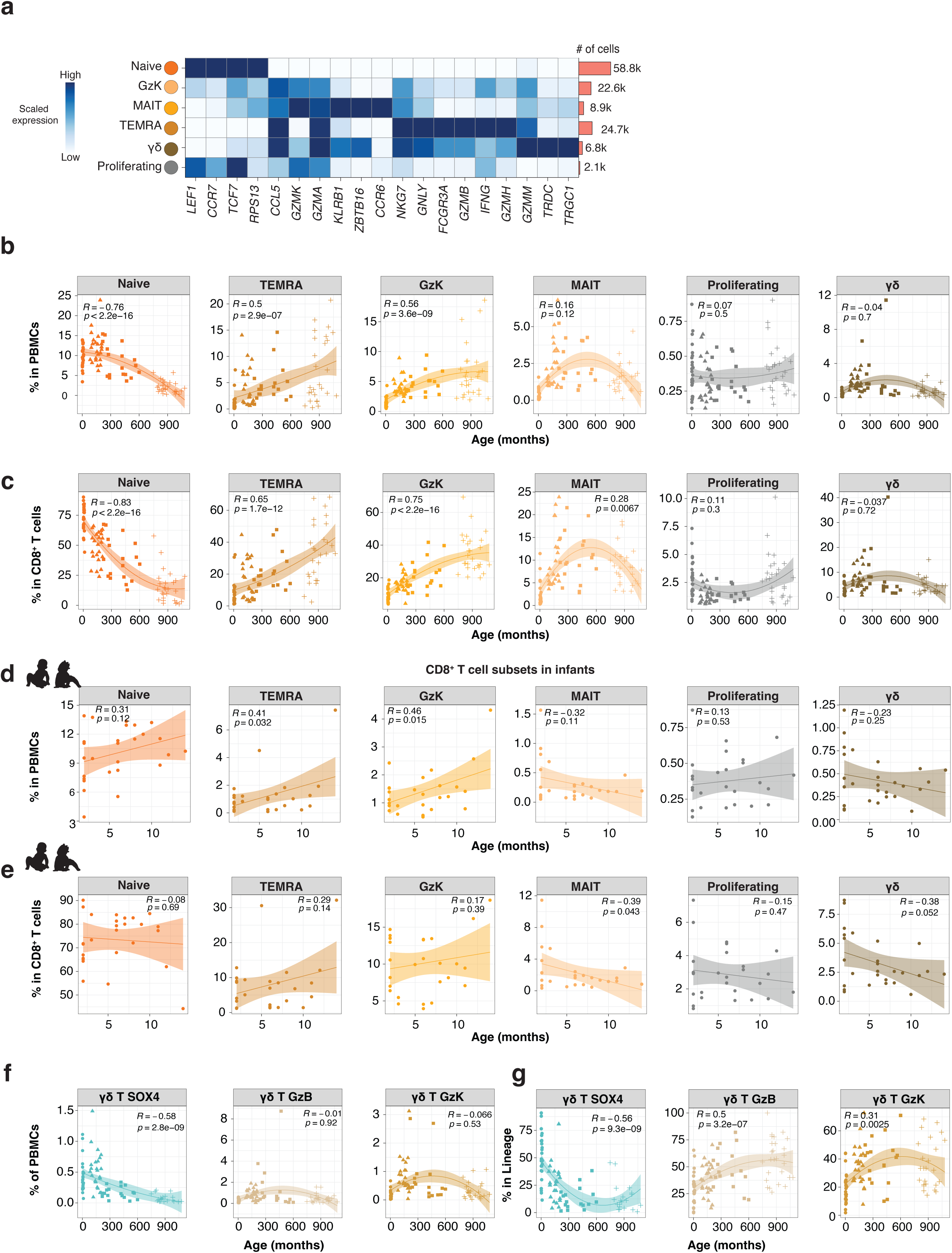
CD8^+^ and γδ T cells-Related to Fig. 6. **a**, Heat map representing scaled expression values of selected marker genes defining each of the CD8^+^ T cell subsets. **b**, Scatterplot representing the frequencies of CD8^+^ T cell subsets in PBMCs (y-axis) and age (x-axis). Each line represents the best-fitted linear regression, with the shading showing the 95% confidence intervals. R represents the Pearson correlation coefficient. **c**, Scatterplot representing the percentage of in each CD8^+^ T cell subsets in total CD8^+^ T cells (y-axis) and age (x-axis). Each line represents the best-fitted linear regression, with the shading showing the 95% confidence intervals. R represents the Pearson correlation coefficient. **d**, As in (**b**) in infant only. **e**, As in (**c**) in infant only. **f**, As in (**b**) in γδ T cell subsets. **g**, As in (c) in γδ T cell subsets.

**Extended Data Fig. 8.**
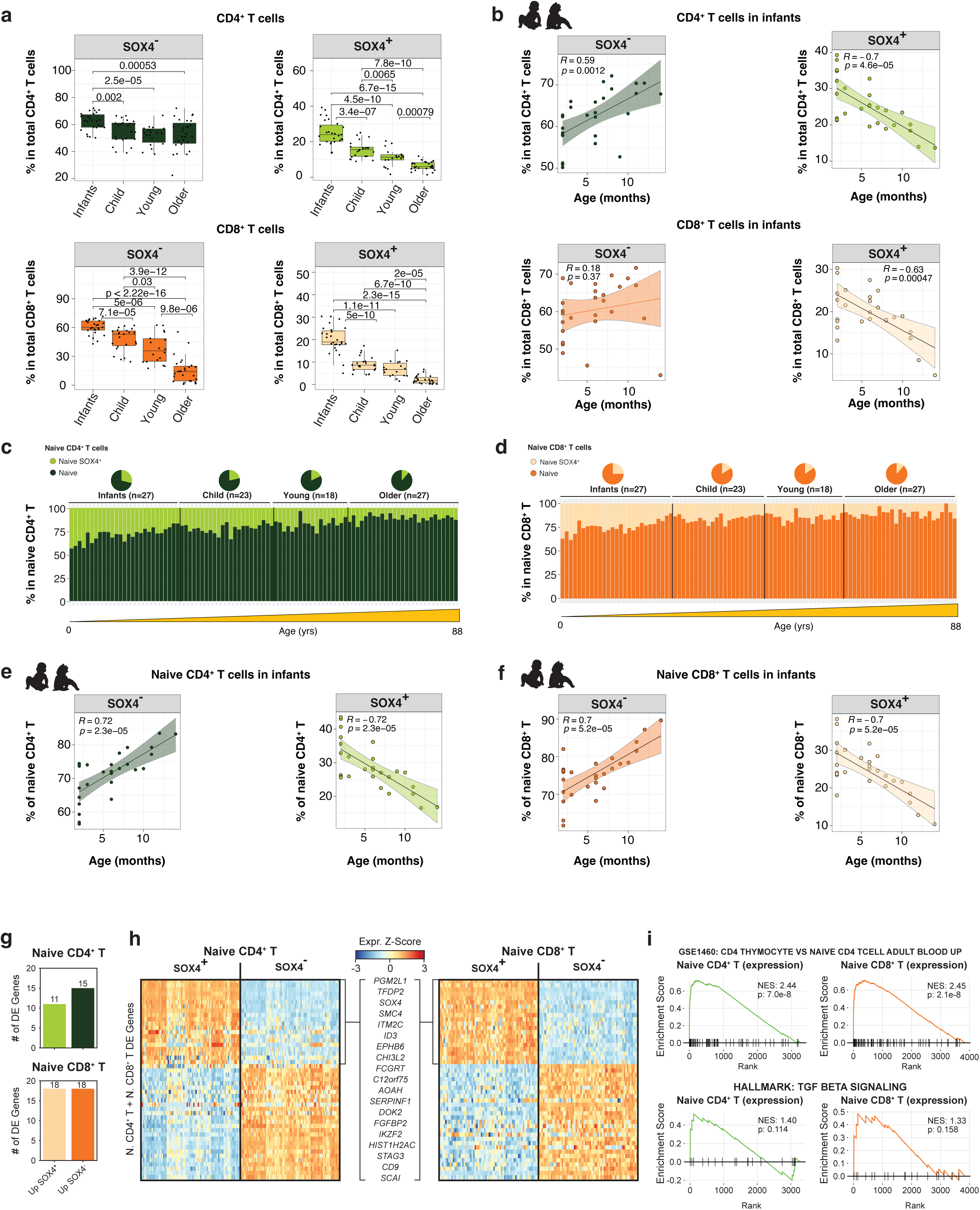
SOX4^+^ T cells (scRNA-seq) – Related to Fig. 7. **a**, Boxplots showing the frequencies of SOX4^+^ CD4^+^ T cells (left panel) and SOX4^+^ CD8^+^ T cells (right panel) in total CD4^+^ T cells or CD8^+^ T cells across the age groups. The upper and lower bounds represent the 75% and 25% percentiles, respectively. **b**, Scatterplot representing the percentage of naïve SOX4^+^ CD4^+^ T cells (left panel) and naïve SOX4^+^ CD8^+^ T cells (right panel) in total CD4^+^ T cells or CD8^+^ T cells (y-axis) and age (x-axis) in infant only. Each line represents the best-fitted linear regression, with the shading showing the 95% confidence intervals. R represents the Pearson correlation coefficient. **c**, Bar plot showing the percentage of SOX4^+^ and SOX4^−^ naïve CD4^+^ T (within total naïve CD4^+^ T cells) across individuals. **d**, Bar plot showing the percentage of SOX4^+^ and SOX4^−^ naïve CD8^+^ T (within total naïve CD8^+^ T cells) across individuals. **e**, Scatterplot representing the percentage of naïve SOX4^+^ (left) and SOX4^+^ (right) in total naïve CD4^+^ T (y-axis) and age (x-axis) in infant only. Each line represents the best-fitted linear regression, with the shading showing the 95% confidence intervals. R represents the Pearson correlation coefficient. **f**, As in (**e**) in naïve CD8^+^ T cells. **g**, Bar plot showing the number of up– and down-regulated differentially expressed (DE) genes in SOX4^+^ (vs. SOX4^−^), in CD4^+^ (upper panel) and CD8^+^ T cells (lower panel). **h**, Heatmap showing scaled expression values of DE genes in SOX4^+^ and SOX4^−^ in total naïve CD4^+^ (left panel) and CD8^+^ T cells (right panel). **i**, Gene enrichment analysis based on DE genes in SOX4^+^ and SOX4^−^ in CD4^+^ (left panel) and CD8^+^ T cells (right panel). Significantly enriched pathways are shown in **Supplementary Table 6**.

**Extended Data Fig. 9.**
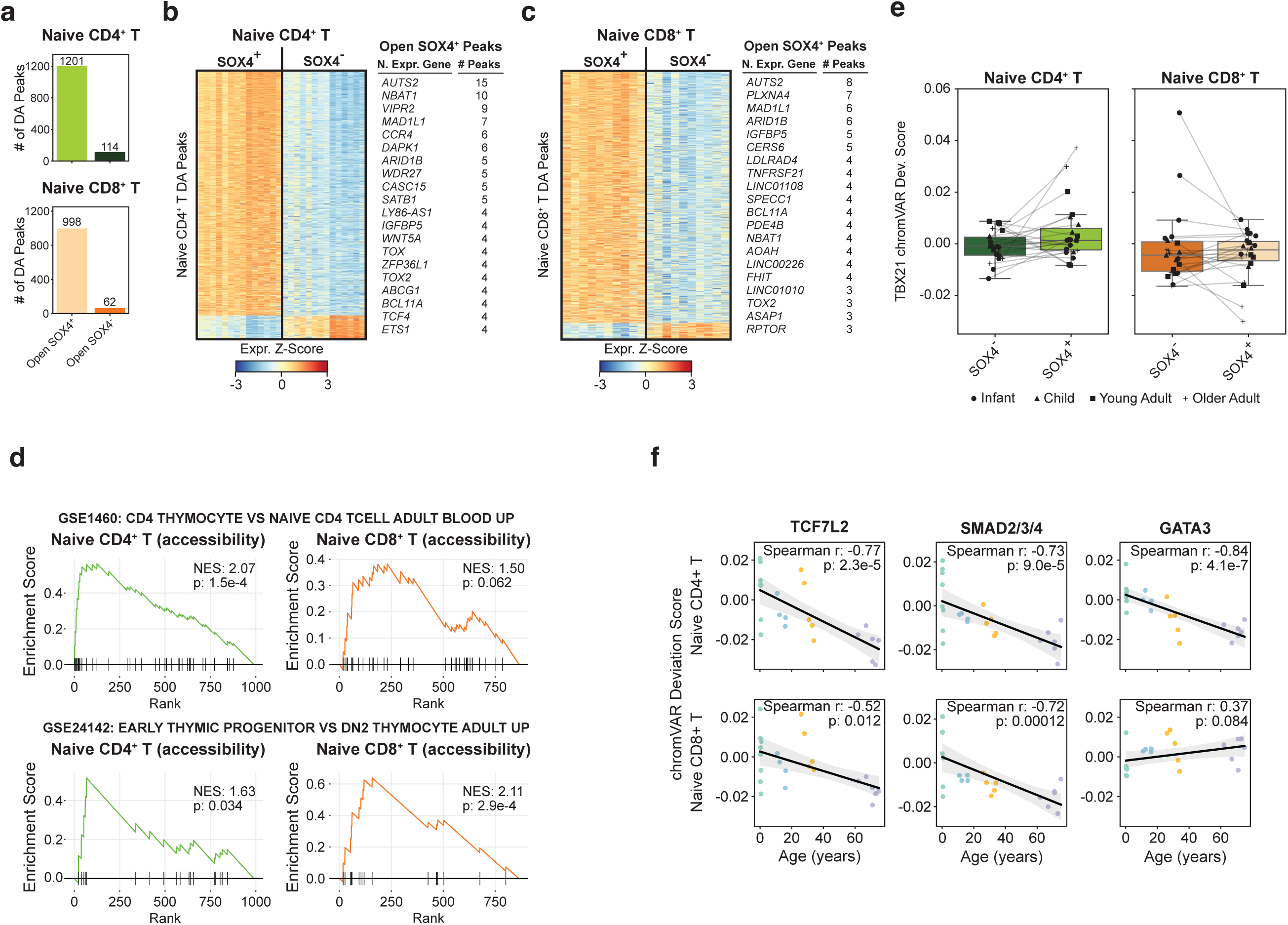
Epigenetic landscape of SOX4^+^ T cells – Related to Fig. 8. **a**, Bar plot showing the number of up– and down-regulated differentially accessible (DA) peaks in SOX4^+^ (vs. SOX4^−^), in total naïve CD4^+^ (upper panel) and CD8^+^ T cells (lower panel). **b**, Heatmap showing scaled expression values of DA peaks in SOX4^+^ and SOX4^−^ in total naïve CD4^+^ T cells. **c**, As in (**b**) in CD8^+^ T cells. **d**, Gene enrichment analysis based on DA peaks in SOX4^+^ and SOX4^−^ in CD4^+^ (left panel) and CD8^+^ T cells (right panel). Significantly enriched pathways are shown in **Supplementary Table 7**. **e**, Box plot representing TBX21 chromVAR deviation scores in SOX4^+^ and SOX4^−^ in CD4^+^ (left panel) and CD8^+^ T cells (right panel). **f**, Scatterplot representing chromVAR deviation scores for TCF7L, SMADs and GATA3 in SOX4^+^ and SOX4^−^ naïve CD4^+^ (upper panels) and CD8^+^ T (lower panels) cells versus age. Each line represents the best-fitted linear regression, with the shading showing the 95% confidence intervals. R represents the Pearson correlation coefficient.

**Extended Data Fig. 10.**
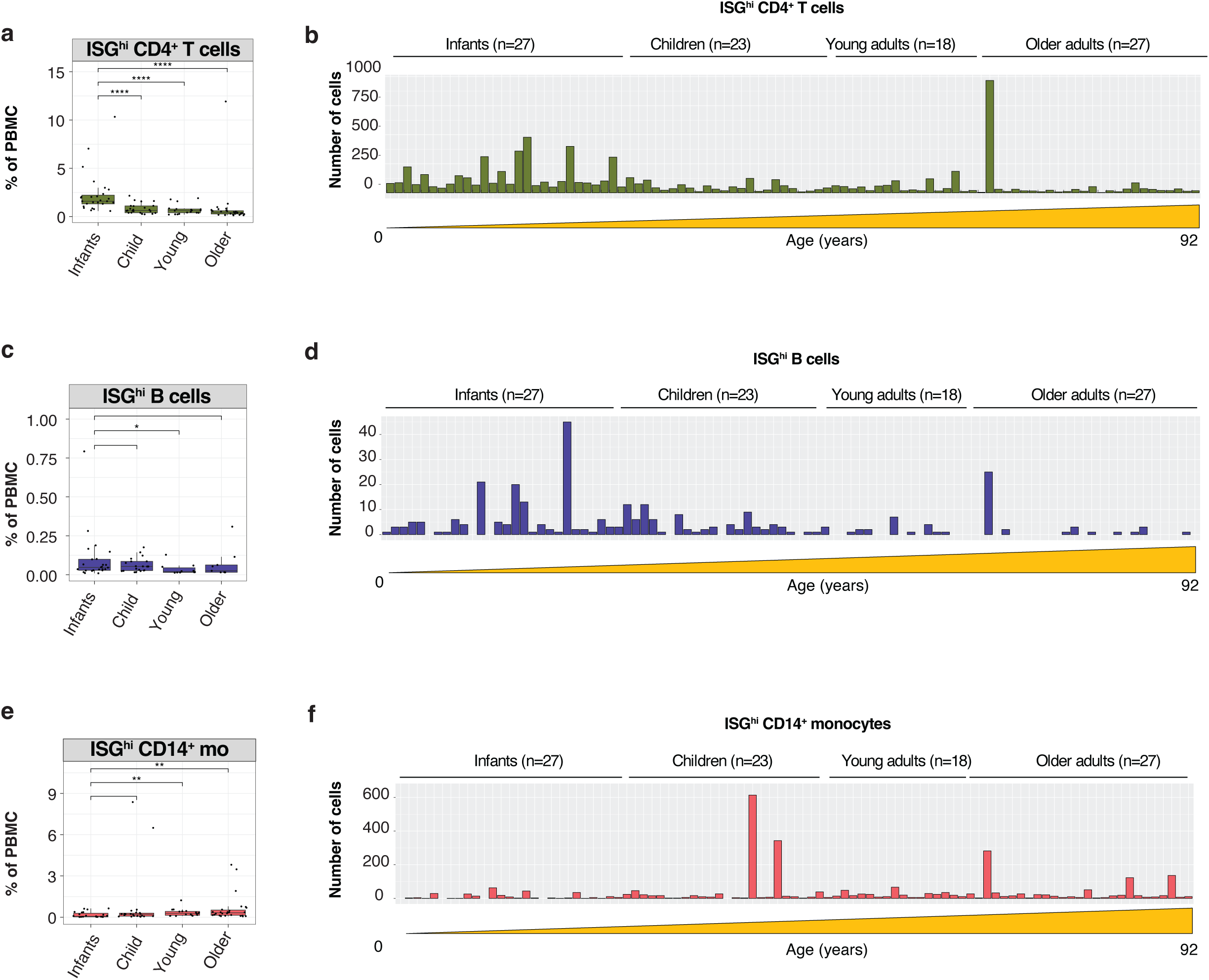
ISG^hi^ subsets. **a**, Boxplots showing the frequencies of ISG^hi^ CD4^+^ T cells (in PBMCs) across the individual (n=95), as categorized into four age groups. *, P<0.05; **, P<0.01; ***, P<0.001: ****, P<0.0001, and ns: non-significant. The upper and lower bounds represent the 75% and 25% percentiles, respectively. **b**, Bar plot showing the number of ISG^hi^ CD4^+^ T cells across individuals (n=95). **c**, As in **(a)** in ISG^hi^ B cells. **d**, As in **(b)** in ISG^hi^ B cells. **e**, As in **(a)** in ISG^hi^ CD14 monocytes. **f**, As in **(b)** in ISG^hi^ B cells.

## Supplementary information

**Supplementary Table 1. Characteristics of enrolled individuals (n=95).**

**Supplementary Table 2. Sequencing information for each sample**

**Supplementary Table 3. Top 100 marker genes in each cluster – related to Fig. 1 and Extended Data Fig. 2**.

**Supplementary Table 4. Top 100 marker genes within each subset.**

**Supplementary Table 5. Subset cell frequencies per donor.**

**Supplementary Table 6. Differentially expressed genes in SOX4^+^ vs. SOX4^−^ T cells.**

**Supplementary Table 7. Differentially accessible peaks in SOX4^+^ vs. SOX4^−^ T cells.**

**Supplementary Table 8. ChromVar results.**

**Supplementary Table 9. SOX4 targets in CD4^+^ and CD8^+^ T cells.**

